# Recapitulating early development of mouse musculoskeletal precursors of the paraxial mesoderm *in vitro*

**DOI:** 10.1101/140574

**Authors:** Jérome Chal, Ziad Al Tanoury, Masayuki Oginuma, Philippe Moncuquet, Bénédicte Gobert, Ayako Miyanari, Olivier Tassy, Getzabel Guevara, Agata Bera, Olga Sumara, Jean-Marie Garnier, Leif Kennedy, Marie Knockaert, Barbara Gayraud-Morel, Shahragim Tajbakhsh, Olivier Pourquié

**Author notes:** Contact Information: Olivier Pourquié, Ph.D., Department of Genetics, Harvard Medical School and Department of Pathology, Brigham and Women’s Hospital, 77 Avenue Louis Pasteur, Boston, MA, USA.

## Abstract

In vertebrates, body skeletal muscles and axial skeleton derive from the paraxial mesoderm which flanks the neural tube and notochord. The paraxial mesoderm forms in the posterior region of the embryo as presomitic mesoderm (PSM), which generates the embryonic segments called somites. Here, we characterized gene signatures identified using microarray series from the mouse PSM and compared the PSM transcriptome dynamics to that of the developing neural tube. In contrast to the PSM where an abrupt transcriptome reorganisation occurs at the level of the determination front, we show that transcriptome changes are progressive during parallel stages of neural tube differentiation. We show that these early differentiation stages of the paraxial mesoderm can be efficiently recapitulated in monolayer culture *in vitro* using murine Embryonic Stem (ES) cells. We describe a serum-containing protocol which parallels *in vivo* tissue maturation allowing differentiation of ES cells towards a paraxial mesoderm fate. We show that R-spondin treatment or Wnt activation alone can induce posterior PSM markers in both mouse and human ES/iPS cells but acquisition of a committed posterior PSM fate requires BMP inhibition to prevent induced cells to drift to a lateral plate mesoderm identity. We show that posterior PSM-like cells induced from mouse ES cells can be further differentiated *in vitro* to acquire an anterior PSM *Pax3*-positive identity. When grafted into injured adult muscle, these induced PSM-like precursors generated large numbers of immature muscle fibers. We further show that exposing ES-derived PSM-like cells to a brief FGF inhibition step followed by culture in horse serum-containing medium allows efficient recapitulation of the myogenic program. Differentiating ES cells first produce mononucleated embryonic myocytes and subsequently multinucleated myotubes, as well as Pax7-positive cells. The protocol described here results in improved differentiation and maturation of mouse muscle fibers differentiated *in vitro* over serum-free protocols. It provides an efficient system for the study of myogenic processes otherwise difficult to study *in vivo* such as fusion or satellite cell differentiation.

## Introduction

Skeletal muscles represent a major derivative of the embryonic paraxial mesoderm. Presomitic mesoderm (PSM) cells, which emerge as a result of gastrulation from the primitive streak and later on the tail bud, express specific sets of genes such as *Mesogenin1* (*Msgn1*) and they experience periodic signalling driven by the segmentation clock while located in the posterior PSM (Hubaud and Pourquie, 2014). Once cells become located in the anterior PSM, at the level of the so-called determination front, they acquire their segmental identity and activate expression of genes such as *Pax3* that control their subsequent differentiation. Epithelial somites form at the anterior tip of the PSM and soon after their formation, distinct subsets of somitic cells begin to activate the myogenic and chondrogenic differentiation programs (Chal and Pourquie, 2009). At the trunk level, the dorso-lateral portion of epithelial somites forms the dermomyotome which contains the Pax3-expressing myogenic precursors. A subset of these precursors located in the dermomyotome lips first activates *Myf5* and then *Myogenin* and gives rise to mononucleated postmitotic myocytes which form the myotome. Precursors of the limb and girdle muscles delaminate from the ventro-lateral lip of the dermomyotome to migrate to their final locations where they activate the myogenic program. Myocytes subsequently fuse in a highly patterned manner to first generate myotubes which further mature into myofibers. Thereafter muscle fibers continue to be formed during embryogenesis from a pool of proliferating precursors expressing Pax3 and Pax7 (Hutcheson et al., 2009). These cells ultimately form the embryonic, fetal and adult muscle fibers and the satellite cells (Biressi et al., 2007).

We recently identified Rspo3 as a secreted protein expressed in the posterior PSM able to induce the differentiation of ES cells toward a Msgn1-positive posterior PSM fate when combined with BMP inhibitors in chemically-defined conditions (Chal et al., 2015). Using chemically-defined culture conditions, these posterior PSM cells can be further induced to differentiate into anterior PSM fates characterized by *Pax3* expression and can be used to subsequently generate large amounts of muscle cells *in vitro* and *in vivo*. Here, we describe a robust protocol based on Wnt signaling activation and BMP inhibition to efficiently produce PSM-like cells from mouse ES cells. We show that the transcriptome of *in vitro* differentiating mouse ES cells exhibits conserved kinetics of gene activation with mouse PSM cells *in vivo*. Furthermore, these induced PSM cells are able to engraft into adult injured muscles, and to generate large amounts of immature skeletal muscle fibers, supporting true paraxial mesoderm commitment. Finally, the PSM precursors could be further differentiated *in vitro* into muscle fibers which are more mature than that generated in serum-free conditions and amenable to long term analysis, thus providing an ideal system to study myogenesis *in vitro* using mouse ES cells.

## Results

### Comparison of the transcriptional landscape between the PSM and posterior Neural Tube in the mouse embryo

We previously reported the generation of a microarray series of consecutive micro-dissected fragments of the E9.5 mouse PSM (Chal et al., 2015). This data set describes the dynamics of the PSM transcriptome from the tail bud to the forming somite. We used this PSM microarray series to identify lists of approximately 40 to 50 highly specific gene signatures for the posterior and anterior PSM domains, respectively (Chal et al., 2015) (Table S1, S2). In addition to genes belonging to pathways well-known to be involved in PSM specification and differentiation such as Wnt, FGF or Notch signaling, the posterior PSM signature was enriched in genes associated to signalling pathways that have been little studied in the context of paraxial mesoderm development (Table S1, S2). Here, we have performed systematic whole mount *in situ* hybridization (ISH) using probes for these genes to validate their expression pattern. Genes identified as strongly expressed in the posterior PSM include the carbohydrate (N-acetylglucosamino) sulfotransferase 7 (*Chst7*), the gene regulated by estrogen in breast cancer product (*Greb1*), Tropomyosin alpha 1 (*Tpm1*), the EGF domain-specific O-linked N-acetylglucosamine (GlcNAc) transferase (*Eogt*), the previously identified cyclic gene *Tnfrsf19 (Troy*), as well as the Apelin receptor (*Aplnr/Apj*) (Kalin et al., 2007) and the Sphingosine phosphate receptors (*S1pr3* and *S1pr5*) (Ohuchi et al., 2008) (Figure 1A, data not shown). Genes expressed in the entire PSM but not in somites included *Greb1L,* the interferon induced transmembrane protein 1 (*Ifitm1, Fragilis2*) (Tanaka and Matsui, 2002). Genes showing expression in the anterior PSM comprised the adducin gamma (Add3), the lipoma HMGIC fusion partner-like 2 (*Lhfpl2*) and Fibulin2 (*Fbn2*) (Figure 1A). A subset of genes was expressed as stripes in the anterior PSM and comprised the transcriptional regulator Myocardin *(Myocd)*, the extracellular matrix protein vitronectin (*Vtn*), the epithelial remodelling factor *Shroom3* (Sousa-Nunes et al., 2003), and a number of genes with unknown function in the PSM including *Abca1, Fam101a (Cfm2)* (Hirano et al., 2005), *Arg1* (Hou et al., 2007), *Pgm5* and *Ism1* (Tamplin et al., 2008) (Figure 1A, data not shown). Thus our data identifies a new set of molecular players showing tightly restricted spatio-temporal expression in the mouse PSM.

**Figure 1:**
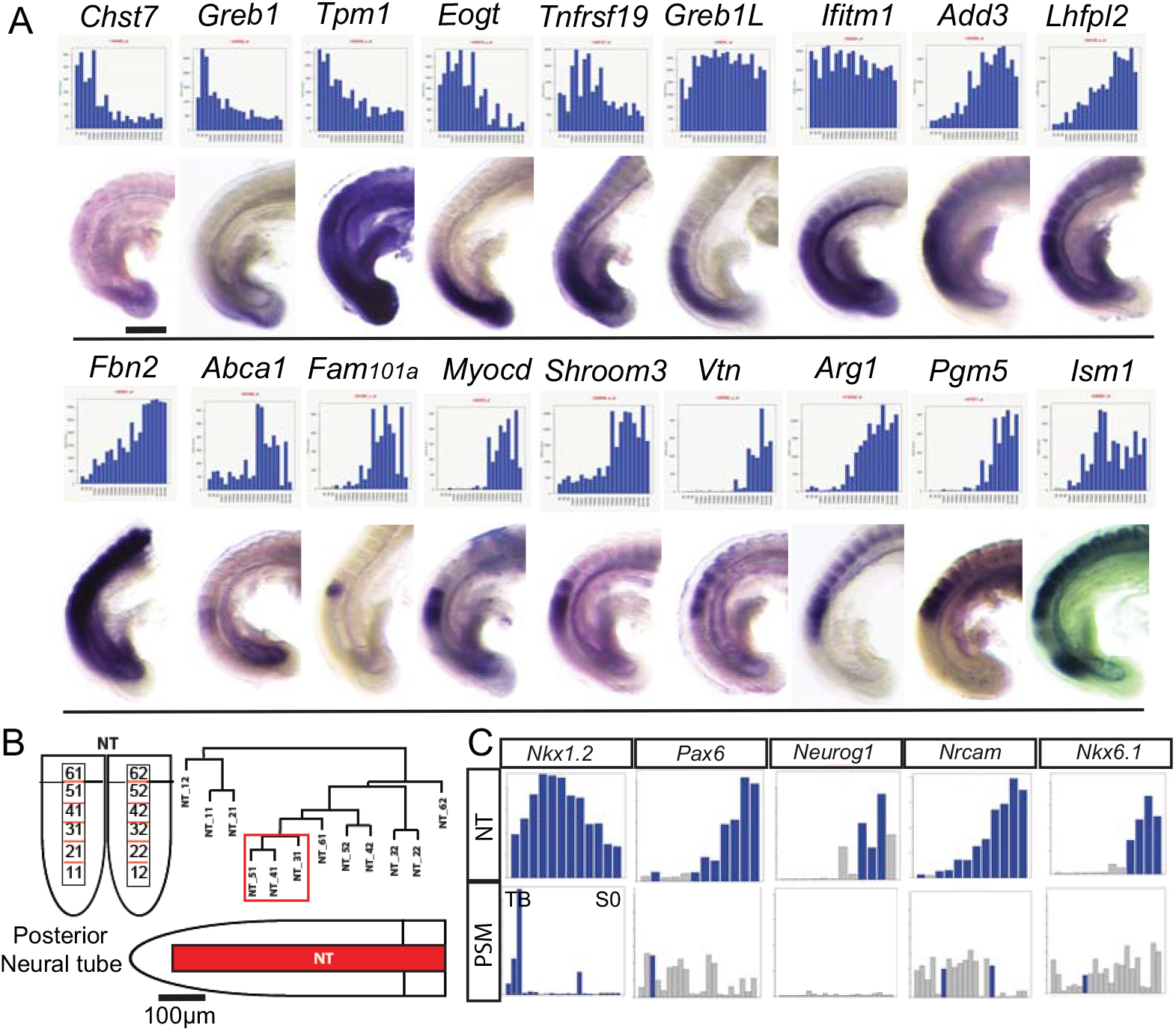
Molecular profiling of early stages of paraxial mesoderm and neural tube differentiation. (A) Validation of novel presomitic mesoderm marker genes identified by the PSM microarray analysis. (top) Expression profiles (blue graphs) of new marker genes as differentially expressed along the PSM microarray series (left, Tail bud domain; right; new somite) (Chal et al., 2015). (bottom) Corresponding validation by *in situ* hybridization on E9.5 mouse embryo tail (lateral view, anterior to the top). Scale bar, 200μm. (B) Hierarchical clustering analysis of the neural tube microarray series. (left) Array series are shown schematically. (right) Series clustering tree, significant clusters are boxed. (bottom) The newly formed neural tube is defined by a single transcriptional domain (red). (C) Validation of the neural tube microarray series. Expression profiles (blue graphs) of key neural genes is compared between the neural tube series (top) and the PSM series (bottom). Each bar corresponds to one array. Note the graded expression in the neural tube (blue) while transcripts levels are very low (grey) in the PSM.

During formation of the posterior body, PSM and neural tube have been shown to share common progenitors in the tail bud, the so-called Neuro-Mesodermal progenitors (NMPs) (Henrique et al., 2015). To evaluate how differentiation programs diverge once cells acquire a neural or paraxial mesoderm fate, we compared the transcriptional program during PSM differentiation with that of the adjacent neural tube. To that end, we generated a series of consecutive microdissected fragments of the posterior neural tube region adjacent to the PSM, from two different E9.5 mouse embryos. Each series comprised six contiguous ~100 μm long neural tube fragments spanning from the tail bud to the level of the newly formed somite (S0) (Figure 1B). Like for the PSM, RNA extracted from each fragment was used to hybridize a single Affymetrix microarray. Analysis of the expression profiles of known genes activated during neural tube differentiation including *Pax6, Neurog1, NrCAM, Nkx6.1* or *Nkx1.2* showed the expected expression gradients in the neural tube (Figure 1C). Thus, the paraxial mesoderm and neural tube microarray series provide an accurate representation of the transcriptional landscape during early stages of the development of these two tissues. Clustering analysis of the PSM microarray series identified an unbiased molecular subdivision of the PSM corresponding to the well-characterized “determination front” at which the segmental prepattern is first established and where differentiation begins (Chal et al., 2015). Such clear demarcation was not observed when a similar analysis was performed for the neural tube series (Figure 1B), suggesting that progressive transcriptional changes accompany early differentiation of the neural lineage (Table S3, S4). Thus our data identify novel PSM specific genes and argue for a different mode of transcriptome regulation during contemporary stages of paraxial mesoderm and neural tube development.

### R-spondin treatment/ Wnt signaling activation in combination with BMP inhibitors promotes posterior PSM differentiation of mouse ES and human iPS cells

*In vivo*, the first stage of paraxial mesoderm cells differentiation corresponds to their exit from the primitive streak or tailbud to enter the posterior PSM. This stage is characterized by the activation of the gene *Mesogenin1* (*Msgn1*), coding for a basic helix-loop-helix transcription factor specifically expressed in the posterior PSM (Yoon et al., 2000). Examination of gene expression profiles of secreted growth factors in the microarray series of differentiating paraxial mesoderm led to the identification of the secreted Wnt agonist *R-spondin3 (Rspo3)*, which is strongly expressed specifically in the posterior PSM (Chal et al., 2015; Kazanskaya et al., 2004). When we treated monolayers of the Msgn1-repV mES reporter cells with 10 ng/mL Rspo3 in a base culture medium containing 15% Foetal Bovine Serum (FBS), we observed a dramatic increase of the percentage of Msgn1-repV^+^ cells over control medium to reach up to 70% after 4 days (Figure 2A). Dimethyl sulfoxide (DMSO) has been shown to promote differentiation of several cell types, including mesoderm from the P19 embryonic carcinoma cell line (McBurney et al., 1982) or from ES cells (Chetty et al., 2013). We also found that addition of DMSO synergized with Rspo3 to induce Venus from Msgn1-repV cells in serum-containing medium or in a serum-free medium (Figure S1, data not shown). Thus, 0.5% DMSO was systematically added to the differentiation medium.

We next tested whether the Venus-positive cells induced by Rspo3 exhibited a characteristic PSM identity. Microarrays from sorted Venus-positive cells cultured for 3 and 4 days in Rspo3 containing medium (RD) were generated and compared the expression of known markers of the PSM. At day 3, many characteristic markers of the posterior PSM such as *Tbx6* were found to be expressed in differentiated ES cells (Figure 2B). Unexpectedly, the *Bmp4* gene, which, in the posterior region of the embryo, is specific for the lateral plate mesoderm, was found to be enriched in the Venus-positive cells differentiated for 4 days in the presence of Rspo3 (Figure 2B). Bmp4 has been shown to promote the lateral plate mesoderm fate at the expense of the paraxial mesoderm and to auto-regulate its own expression (Adelman et al., 2002; Tonegawa et al., 1997). Accordingly, Msgn1-repV^+^ cells induced in RD media also expressed the lateral plate marker *Foxf1a* (Figure 2B). The expression of lateral plate markers in the cell population expressing *Msgn1* suggests that treatment with Rspo3 alone leads cells to acquire a mixed identity between paraxial mesoderm and lateral plate.

**Figure 2:**
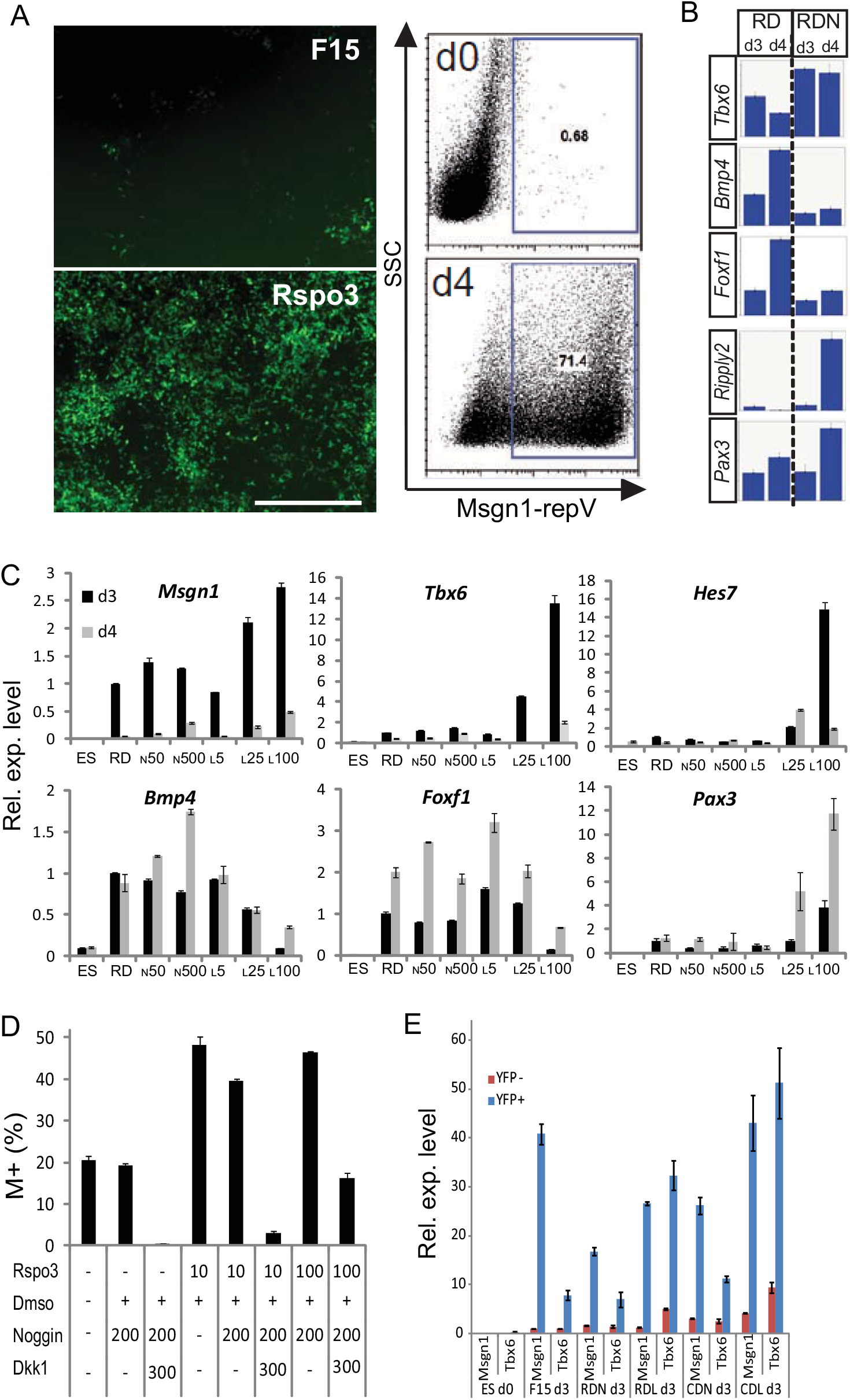
Wnt pathway activation and BMP pathway inhibition are required to induce a PSM-like fate from mouse ES cells. (A) Activation of Msgn1-repV reporter with R-spondin3 treatment. (left) Detection of the fluorescent reporter activation after 4 days of differentiation of Msgn1-repV ES cells, in control 15% FBS (F15) containing medium or supplemented with recombinant Rspo3 (10ng/mL). Scale bar, 200μm. (right) Representative flow-cytometry analysis of Msgn1-repV cultures at day 0 (d0) and day 4 (d4) of differentiation in F15 medium supplemented with Rspo3 at 10ng/mL, showing the induction of more than 70% of Venus (YFP) -positive cells. SSC: Side Scatter. (B) Comparison of *Tbx6, Bmp4 Foxf1, Ripply2 and Pax3* expression levels as detected in microarrays from FACS -sorted Msgn1-repV^+^ cells cultured for 3 (d3) and 4 (d4) days in medium containing Rspo3 and DMSO (RD) and in medium containing Rspo3, DMSO and Noggin (RDN). (C) qPCR analysis of *Msgn1, Tbx6, Hes7, Bmp4, Foxf1* and *Pax3* expression levels comparing undifferentiated Msgn1-repV cells (ES) with FACS -sorted Msgn1-repV^+^ cells cultured for 3 (black bars) or 4 days (grey bars) in Rspo3, DMSO (RD) medium containing 0, 50 (N50), 500 (N500) ng/mL of Noggin or 5 (L5), 25 (L25), 100 (L100) nM of the Ldn BMP inhibitor. Values are normalized to RD d3 condition. Mean +/- s.d. (D) qPCR analysis of *Msgn1* and *Tbx6* expression levels comparing Msgn1-RepV -negative (YFP-, red) and -positive (YFP+, blue) cell fractions sorted at day3 of differentiation in control FBS15% serum or containing Rspo3 (R), Chir (C), Dmso (D), Noggin (N) and Ldn (L). Expression level in ES cells is also shown. Expression are normalized to F15 d3 values. Mean +/- s.d. (E) Impact of Wnt inhibition on Msgn1-RepV induction. Dose response of the effect of Rspo3 on Msgn1-RepV^+^ (M^+^) cells induction after 4 days of differentiation in the presence or absence of the different concentrations of the Wnt inhibitor Dkk1 in medium containing 15% FBS, DMSO and Noggin. Concentrations in ng/mL. Mean +/- s.d.

In order to prevent cells to acquire a lateral plate identity, we added the BMP inhibitor Noggin to the culture medium. Microarray and qPCR analysis indicated that Noggin-treated cells down-regulated *Bmp4* and *Foxf1a* and upregulated *Tbx6* (Figure 2B). Furthermore, after 4 days in culture, Noggin-treated cells up-regulated the anterior PSM-specific markers *Ripply2* and *Pax3* (Figure 2B). Stronger inhibition of Bmp4 activity was observed with the chemical BMP inhibitor LDN 193189 (thereafter named Ldn), a more potent derivative of dorsomorphin (Cuny et al, 2008). qPCR analysis on FACS-sorted Msgn1-repV^+^ cells revealed that Ldn applied together with Rspo3 on mouse ES Msgn1-repV cells acts in a dose-dependent fashion, inhibiting *Foxf1a* and *Bmp4* expression and promoting expression of *Tbx6, Hes7, Msgn1* and *Pax3* much more efficiently than Noggin (Figure 2C). Addition of Noggin or Ldn to the medium was however found to lead to lower levels of induction of Venus^+^ cells from the Msgn1-repV cultures (data not shown). Substituting Rspo3 by Rspo2 also led to similar results (Figure S1). Together, our data demonstrates that efficient differentiation of mouse ES cells toward a posterior PSM fate can be achieved by culturing the cells in a medium containing Rspo3, DMSO and Ldn (thereafter named RDL).

We next studied the mechanism of action of Rspo3. R-spondins are secreted molecules that bind Lgr4 and Lgr5 receptors and activate both the canonical and planar cell polarity (PCP) Wnt signalling pathways (de Lau et al., 2012). While *Lgr5* expression was not significantly detected in the PSM in our profiling series, expression of *Lgr4* was observed both in the PSM and neural tube ((Chal et al., 2015) and data not shown). We next asked whether Rspo3 activates canonical Wnt signaling in differentiating mouse ES cells. Addition of Dkk1 inhibited the induction of Venus^+^ cells from Msgn1-repV cells by Rspo3 (Figure 2D). Furthermore, substituting Rspo3 by the GSK3-inhibitor and canonical Wnt activator CHIRON 99021 (Chir) led to comparable induction level of the Msgn1-repV^+^ population (Figure 2E, Table S5 and data not shown). These data demonstrate that Rspo3 efficiently promotes the induction of a Msgn1-positive cell population by activating the canonical Wnt pathway, supporting an important role for this pathway in PSM specification.

We next analyzed induction of the posterior PSM fate from human iPS cells *in vitro.* To monitor the differentiation of hiPS cells toward paraxial mesoderm, we generated a fluorescent reporter line harboring a MSGN1-Venus fusion transcript by knock-in using the CRISPR/Cas9 system (Figure 3A). hMSGN1 reporter cells were differentiated in serum-free medium containing Chir/Ldn (CL) for 4-5 days to induce paraxial mesoderm differentiation (Chal et al., 2016; Chal et al., 2015). Venus was expressed in induced cells starting at day 3 and the *MSGN1* transcript was almost exclusively found in the Venus^+^ (hM^+^) fraction (Figure 3B). By 4 days of differentiation, flow cytometry analysis demonstrates that up to 95% of the differentiating hiPS cells were Venus-positive, both in Chir/Ldn (CL) and in Chir only (C) conditions (Figure 3C). hM^+^ cells induced in the presence of Ldn also expressed the PSM-specific marker TBX6, validating the specificity of the reporter activity and the efficiency of the protocol to induce posterior PSM fate in human cells (Figure 3D). Next, we compared the gene expression profile of FACS-sorted hM^+^ cells generated in serum-free Chir/Ldn (CL) containing medium to undifferentiated hPSC (Figure 3E, Table S6). Mouse posterior PSM signature genes including *DKK1, MSGN1, RSPO3, BRACHURY, HES7* and *LFNG* were strongly up-regulated in the differentiated hM^+^ cells (Figure 3E). To evaluate the impact of BMP inhibition on the gene signature of induced hM^+^ cells, microarrays from FACS-sorted hM^+^ cells cultured in serum-free Chir/Ldn (CL) or Chir only (C) medium were compared (Figure 3F). Genes of the Wnt (*AXIN, DKK1, DKK2, RSPO3, LEF1, WNT3A*), Notch (*DLL1, DLL3, LFNG, HES7*) and FGF (*DUSP1, FGF10, FGF8, FGF9*) signaling pathways which are important for posterior PSM development were found upregulated in hM^+^ cells induced in CL conditions (Fold change >2 and False discovery rate <10%). By contrast, hM^+^ cells induced in the presence of the WNT activator alone up-regulated lateral plate and cardiac markers including *FOXF1, BMP4, TBX3, HAND1, HAND2, CXCR7* while posterior PSM markers were expressed at a lower level compared to cells differentiated in CL (Figure 3F, G, Table S7). Strikingly, the anterior paraxial mesoderm marker *Pax3* was found to be up-regulated in absence of LDN (Figure 3F). Together these data suggest that these cells exhibit a mixed paraxial/lateral plate mesoderm identity. Thus, like for mouse ES cells, efficient induction of the paraxial mesoderm fate from human iPS cells requires BMP inhibition at the critical stage of posterior PSM differentiation, preventing early paraxial mesoderm cells from drifting to a lateral plate mesoderm fate.

**Figure 3:**
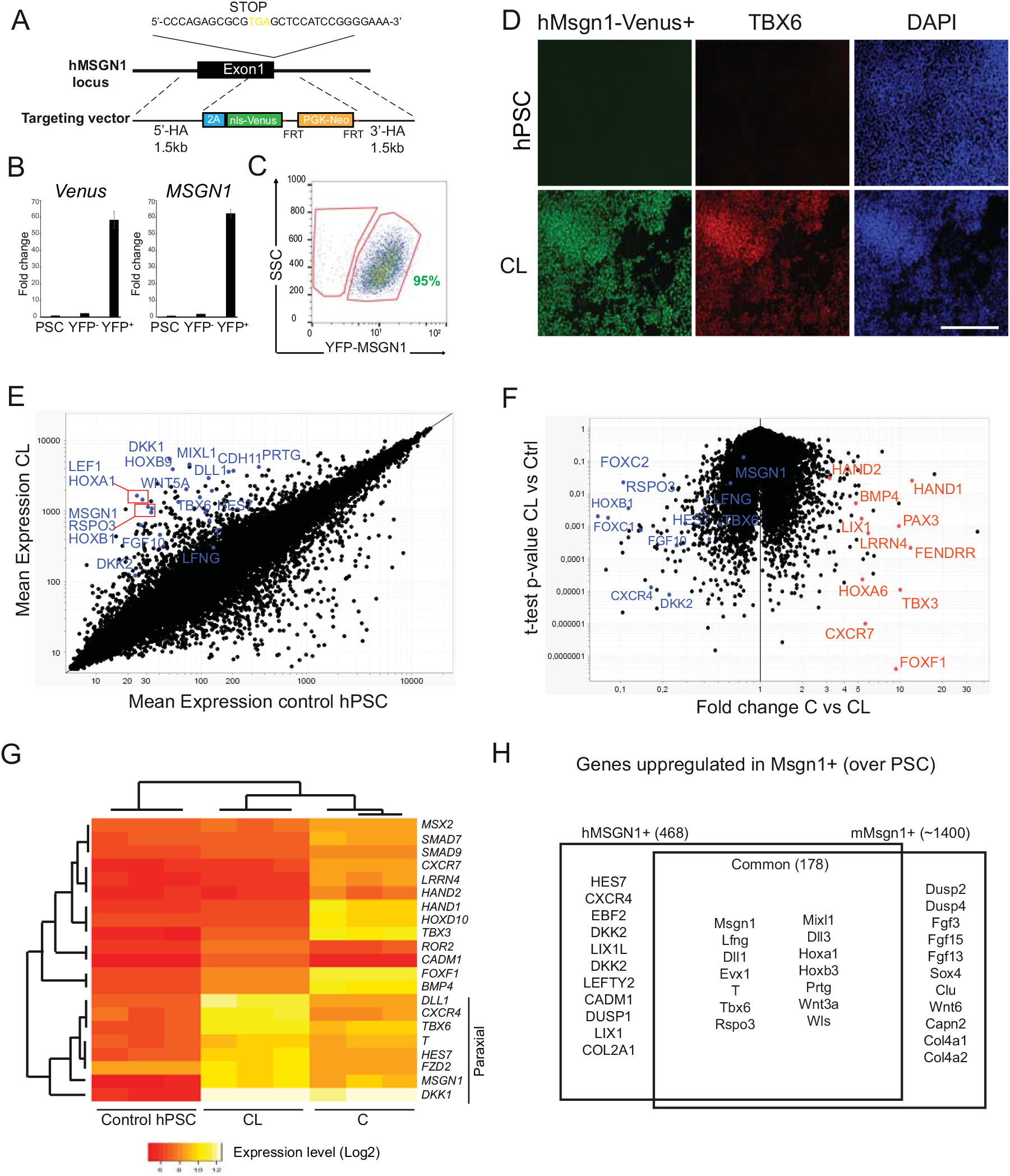
Generation of hMSGN1^+^ PSM-like progenitors from hPSC. (A) Targeting strategy for hMSGN1-Venus reporter generation with CRISPR/Cas9. HA: homology arm (B) Expression of *Venus* and *MSGN1* transcript in FACS-sorted hMSGN1-Venus^+^ (YFP+) compared to Venus^-^ (YFP-) and undifferentiated hPSCs, differentiated for 4–5 days in serum-free medium containing Chir/Ldn according to (Chal et al., 2016; Chal et al., 2015). Values are normalized to hPSC condition. Mean +/- s.d. (C) Representative flow cytometry analysis of hMSGN1-Venus culture differentiated for 4 days in serum-free medium containing Chir/Ldn. SSC: Side scatter (D) Expression of the hMSGN1-Venus reporter after 4 days of differentiation in serum-free medium containing Chir/Ldn (CL medium) and stained with an anti-TBX6 antibody and Dapi. Undifferentiated hPSCs are also shown as negative control. (E) Comparison of gene expression profiles of FACS-sorted hM^+^ cells generated in serum-free Chir/Ldn (CL) to undifferentiated hPSC. The mean expression for each probe set (black dot, logarithmic scale) is shown. Most posterior PSM signature genes found in differentiating mouse ES were also found upregulated in differentiated hM^+^ cells (blue). (F) Comparison of the gene expression profiles of FACS-sorted hM^+^ cells differentiated for 4 days in serum-free Chir/Ldn (CL) or CHIR only (C) medium. Volcano plot showing the fold change expression for each probe set (black dot, logarithmic scale) and the associated p-value. Representative marker genes are highlighted for each condition. (blue) Chir/Ldn enriched genes, (orange) Chir-only enriched genes. (G) Expression heatmap for lateral plate and presomitic (paraxial) mesoderm marker genes comparing hMSGN1-Venus -positive cells differentiated for 4 days in serum-free Chir/Ldn (CL) or Chir-only (C) medium, and to undifferentiated hPSC. (H) Venn diagram comparing the genes upregulated in mouse and human Msgn1-Venus^+^ following normalization to undifferentiated mESC and hiPSC, respectively. For each category, representative marker genes are shown. hM^+^ and their mouse counterpart share the Core posterior PSM signature.

We next compared hM^+^ progenitors induced *in vitro* by Wnt activation and BMP inhibition to their mouse counterparts (Chal et al., 2015). As observed for the mouse Msgn1-repV^+^ differentiated in serum-free CL conditions, hM^+^ cells upregulated a large number of posterior PSM signature genes, including *MSGN1, TBX6, BRACHURY, WNT5A, RSPO3, CDX2*, and *EVX1* (Figure 3H; Table S5, S6). Interestingly, a subset of genes were found differentially enriched specifically in human versus mouse M^+^ progenitors. hM^+^ progenitors expressed genes such as *DKK2* and *CXCR4* which were detected at significantly lower level in mouse M^+^ cells, while mouse M^+^ cells show enriched expression of some FGF pathway components (*Dusp2, Dusp4, Fgf3, Fgf15*) when compared to their human counterparts. Altogether, our data indicate that as reported for mouse ES cells, *in vitro* differentiation of human posterior PSM-like cells can be induced in the presence of a WNT activator and a BMP inhibitor. Moreover, our data suggest that the transcriptome of mouse and human posterior PSM-like cells is highly similar.

### Generation of Pax3-positive anterior PSM precursors from ES cells *in vitro*

*In vivo,* as paraxial mesoderm cells mature, they become located in the anterior PSM, and down-regulate the expression of *Msgn1* while activating the expression of *Pax3* at the level of the determination front. FACS-sorted mouse M^+^ cells differentiated in RDL or CDL for 3 days strongly express the posterior PSM markers *Tbx6* and *Hes7* compared to M^-^ cells, indicating that they share features with posterior PSM cells (Figure 2D, 4A). After 4 days of culture in RDL M^+^ cells activated anterior PSM and early somite markers including *Pax3, Meox1/2, Paraxis/Tcf15, Foxc1/2, Tbx18*, and *Uncx4.1* (Figure 4A-C). While *in vivo*, these genes are expressed in a more anterior PSM domain than *Msgn1*, their co-expression with the Venus protein is due to the stability of the reporter which persists for some time in cells that have ceased expressing *Msgn1*. M^+^ cells induced in basal (FBS 15%, F15) medium without Ldn strongly up-regulated BMP4 at day 4-5 (Figure 4A-C). This transition to an anterior PSM fate was highly dependent on the concentration of Ldn, with 100nM leading to the highest induction of anterior PSM markers, and the lowest expression levels of the lateral plate markers (Figure 4B). Chir (CDL medium) was found to be as efficient as R-spondin3 at inducing the anterior PSM fate (Figure 4C) while cells differentiated in base F15 medium failed to activate the anterior PSM program. Therefore, our data suggest that mouse ES cells differentiated in RDL or CDL media are able to efficiently differentiate into anterior PSM and to activate the early somitic differentiation program.

**Figure 4:**
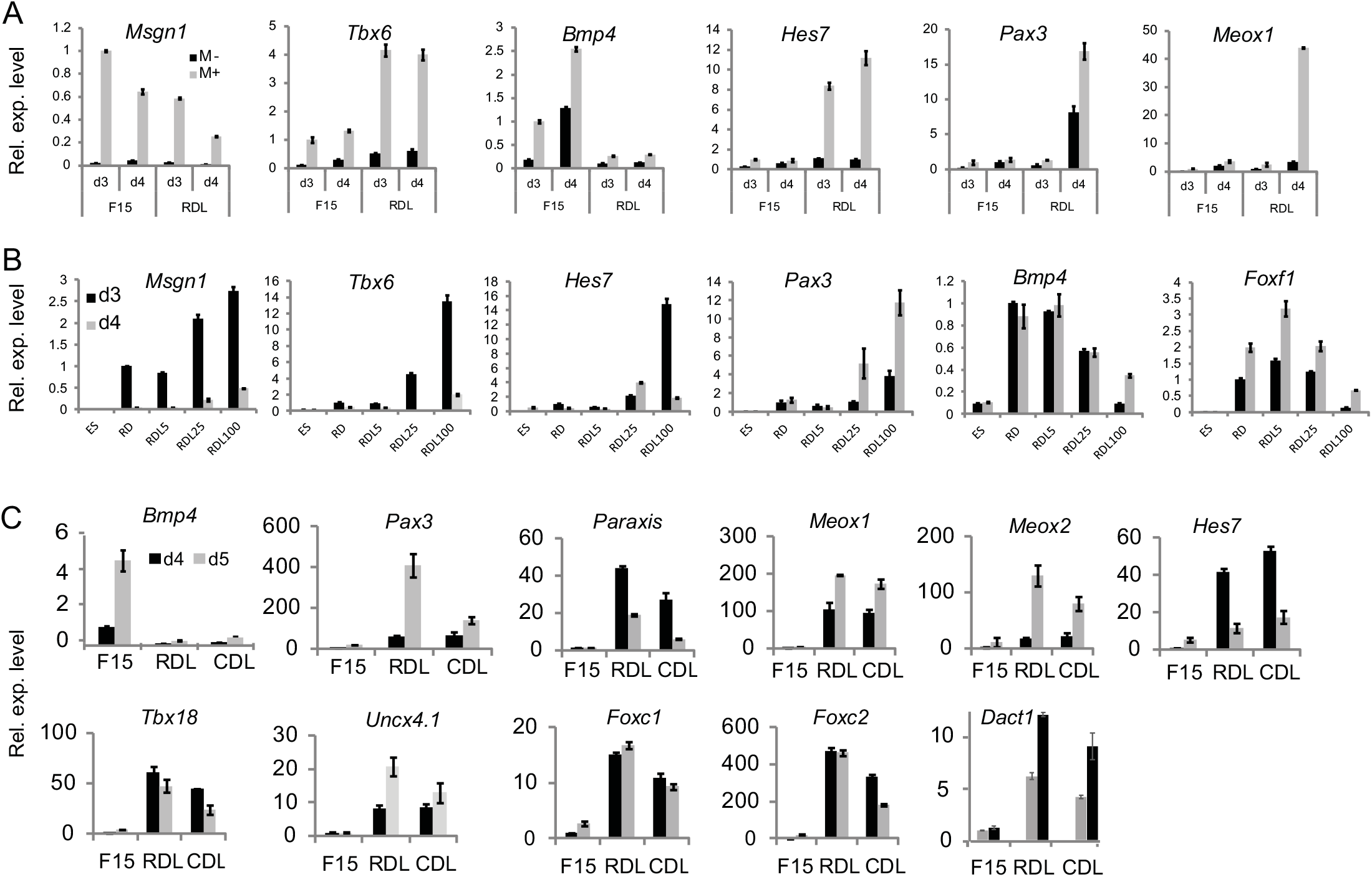
Differentiation of *Pax3^+^* PSM-like progenitors from ES cells *in vitro*. (A) qPCR analysis comparing the expression level of *Msgn1, Tbx6, Bmp4, Hes7, Pax3*, and *Meox1* in FACS-sorted Msgn1-repV -positive (M^+^, grey bars) and -negative (M-, black bars) cells cultured for 3 (d3) or 4 (d4) days in 15% FBS (F15) or Rspo3/DMSO/Ldn (RDL) medium. Values are normalized to the M^+^ F15 d3 condition. Mean+/- s.d. (B) Dose-response analysis of the effect of Ldn concentration on differentiation of Msgn1-repV^+^ cells to a PSM fate. qPCR expression level of *Msgn1, Tbx6, Hes7, Pax3, Bmp4*, and *Foxf1* in FACS-sorted Msgn1-repV^+^ on day 3 (d3) or 4 (d4) of differentiation. Note the transition from a Msgn1^+^ stage (posterior PSM) to a Pax3^+^ stage (anterior PSM). Values are normalized to the RD d3 condition. Mean+/- s.d. (C) Comparison of the anterior PSM /somitic markers induction in FACS-sorted Msgn1-repV^+^ progenitors in presence of Rspo3 (RDL) or Chir (CDL), in media containing DMSO and Ldn. Values are normalized to the F15 d4 condition. Mean+/- s.d.

In order to better monitor the differentiation efficiency of mouse ES cells into anterior PSM precursors, we took advantage of the mouse ES Pax3-GFP reporter line (Chal et al., 2015). During development, *Pax3* is expressed both in mesodermal and neural derivatives, but fluorescence intensity of the reporter was found to be weaker in the former. When these cells were differentiated in F15 or in R-spondin3-containing medium without Ldn (RD), about 1% of GFP^+^ cells was detected after 5 days in culture (Figure 5A). In contrast, in RDL medium, up to 20-40% of GFP^+^ cells were observed at day 5 (Figure 5A). The peak of Pax3-GFP expression followed by approximately one to two days the peak of Msgn1-repV expression. Maximal induction of the Pax3-GFP^+^ cells was observed when Ldn was added in the differentiation media at day 0 (Figure 5B). We confirmed that at day 5-6, Pax3-GFP^+^ cells in culture also expressed the Pax3 protein (Figure 5C). Pax3 is expressed both in the paraxial mesoderm and in neural precursors. However, the Pax3-GFP^+^ cells induced in our RDL/ CDL conditions exhibit a characteristic mesenchymal aspect, distinct from the rosette-forming Pax3^+^ neural precursors found in neural-inducing (dual Smad inhibition) conditions (Figure 5D) (Chambers et al., 2009). The Pax3-GFP^+^ cells were also found to be essentially negative for the neural marker *Sox2* but expressed the anterior PSM/somitic markers *Pax3, Uncx, Meox1 and Foxc2* at levels comparable to those detected *in vivo* (Figure 5E). Thus, differentiation of ES cells in RDL or CDL media was found to efficiently recapitulate the major stages of PSM differentiation *in vitro*.

**Figure 5:**
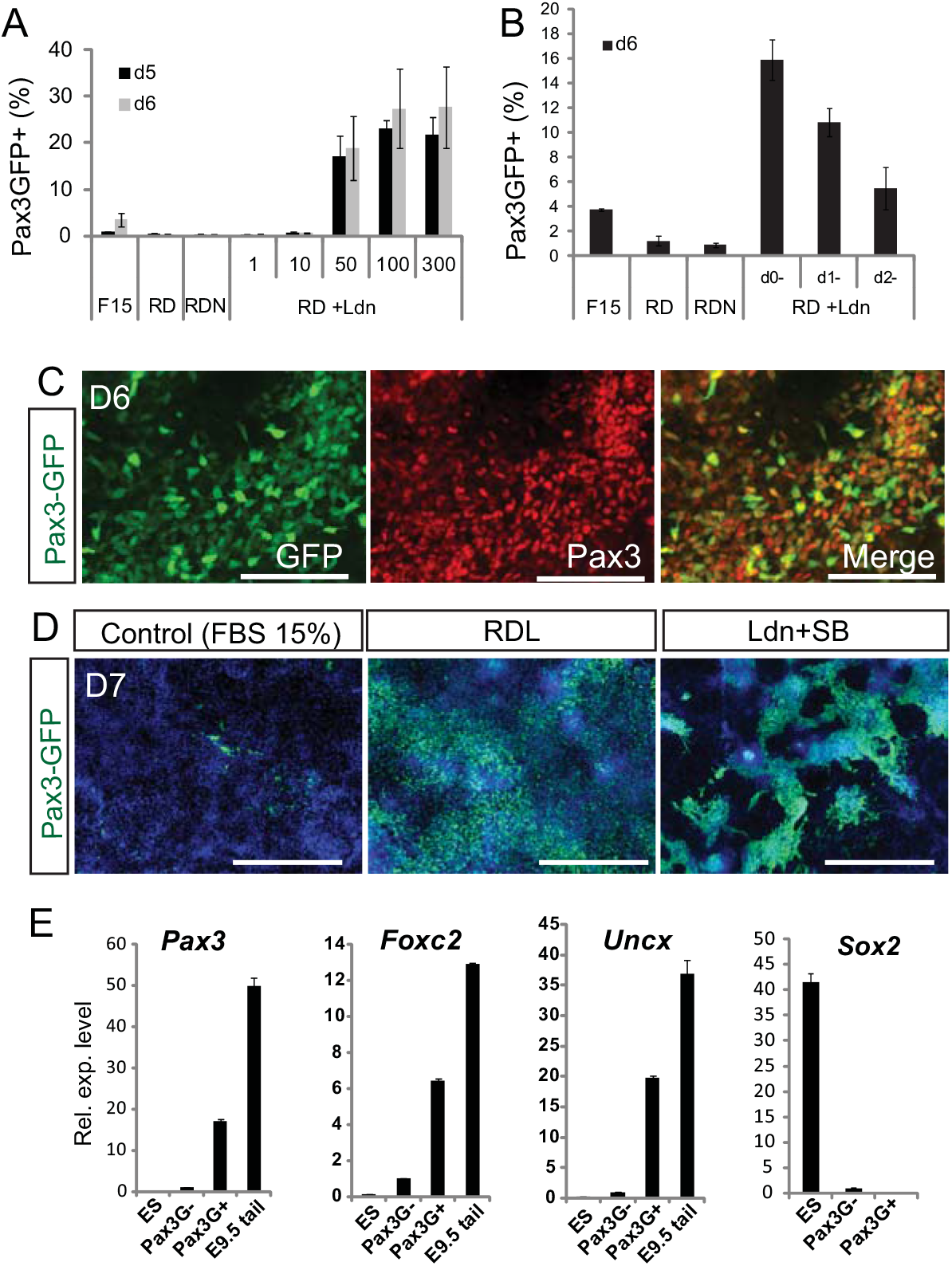
Generation of Pax3-GFP^+^ cells exhibiting anterior PSM identity *in vitro*. (A) Induction kinetics of the Pax3-GFP reporter in mouse ES cell culture differentiated for 5 (d5) or 6 (d6) days in control 15% FBS (F15) medium, in Rspo3-containing medium alone (RD) or supplemented with Noggin (RDN) or with a range of Ldn concentration (+Ldn, in nM). Note robust induction of Pax3-GFP for Ldn concentration above 50nM. Mean+/- s.d. (B) Analysis of the temporal requirement for Ldn for Pax3-GFP^+^ cells generation. Induction kinetics of the Pax3-GFP reporter in mouse ES cultures differentiated for 6 (d6) days in control 15% FBS (F15) medium, in Rspo3 -containing medium alone (RD) or supplemented with Noggin (RDN) or with a 0.1μM Ldn supplemented at day 0, 1 or 2 of differentiation. Mean+/- sd (C) Representative GFP (left, green) and Pax3 immunostaining (center, red) on differentiated cultures of mouse Pax3-GFP ES cells differentiated in RDL medium for 6 days. Scale bar, 100μm. (D) GFP immunostaining on differentiated cultures of mouse Pax3-GFP ES cells differentiated in control 15% FBS (F15) medium, RDL medium or neurogenic (dual smad inhibition, SB431542 +Ldn) medium. Note the distinct morphology of the GFP^+^ cells in RDL versus neurogenic conditions. Scale bar, 400μm. (E) qPCR analysis comparing the expression level of the genes *Pax3, Foxc2, Uncx* and *Sox2* expression levels in undifferentiated Pax3-GFP cells (ES), in FACS-sorted Pax3-GFP -positive (Pax3G+) and negative (Pax3G-) cells after 6 days of differentiation in RDL medium. Expression levels in E9.5 mouse embryonic tail is also provided for comparison. Values are normalized to the Pax3G- condition. Mean+/- s.d.

### Validation of the paraxial mesoderm identity of mouse ES cells differentiated *in vitro*

The presence of PDGFRα (CD140a) and the absence of VEGFR2 (also named Flk1, KDR or CD309) have been used to identify the paraxial mesoderm lineage in differentiated ES cells *in vitro* (Darabi et al., 2008; Nishikawa et al., 1998; Sakurai et al., 2006; Sakurai et al., 2009). *In vivo*, however, these surface markers are not specific to the paraxial mesoderm and their expression largely overlaps with other mesodermal populations such as the lateral plate (Ding et al., 2013; Ema et al., 2006; Motoike et al., 2003). FACS analysis of the Msgn1-repV cultures differentiated for 3-4 days in RDL medium with or without a pre-differentiation step in N2B27 medium supplemented with 1% Knock-out serum (thereafter, NK1) medium (Chal et al, 2015), revealed that more than 90% of the Msgn1-repV^+^ (M^+^) cells are recognized by the PDGFRα antibody (Figure 6A). However, 25% of these M^+^ cells were also found to express VEGFR2 (Figure 6B). Moreover, PDGFRα was not specific to the Msgn1-repV^+^ population as it also marked 75% of the Msgn1-repV^-^ (M^-^) cells (Figure 6B), 45% of which also expressed VEGFR2 (Figure 6A, B). We also analyzed expression of the surface marker CXCR4 (Borchin et al., 2013) in relation to M^+^ cells and found that while most of the M^-^ were CXCR4+, about 30-50% of the M^+^ cells were also CXCR4^+^, suggesting the idea that CXCR4 cannot discriminate for PSM fate *in vitro* (Figure S2).

**Figure 6:**
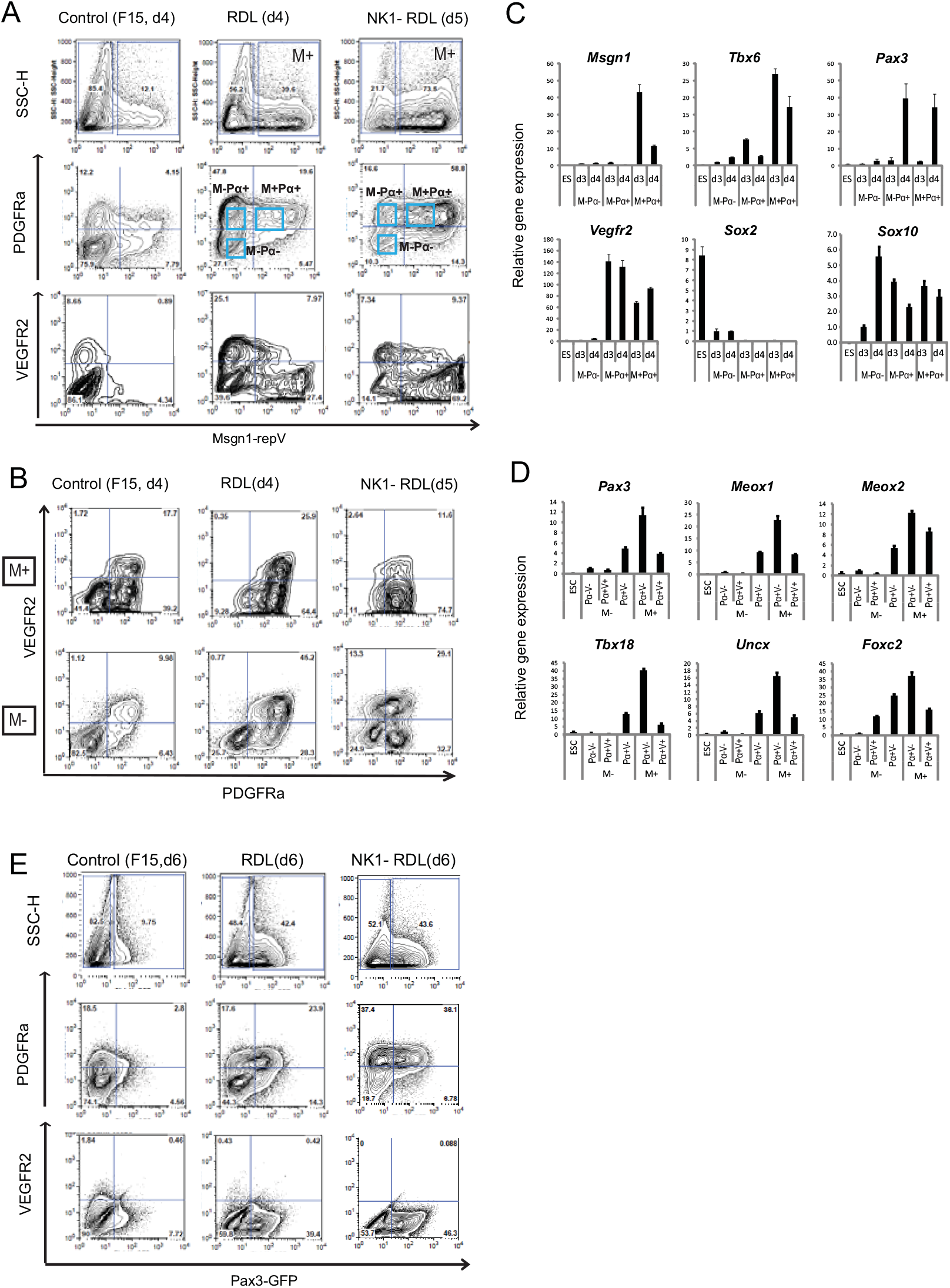
Analysis of PDGFRα and VEGFR2 expression in mouse ES-derived PSM-like cells. (A) Representative FACS analysis of Msgn1-repV reporter cell cultures differentiated *in vitro* in (left) control FBS 15% (F15), in RDL medium without (center) or with pre-differentiation in NK1 medium (NK1- RDL, right) for 4-5 days and labeled with anti-PDGFRα or anti-VEGFR2 antibodies. Distinct subpopulations are shown (blue boxes). Pα: PDGFRα. (B) Representative analysis of PDGFRα and VEGFR2 expression in the Msgn1-RepV- positive (M^+^) and -negative (M-) populations, differentiated in (left) control FBS 15% (F15), in RDL medium without (center) or with pre-differentiation in NK1 medium (NK1- RDL, right) for 4-5 days. (C) qPCR analysis of the expression of *Msgn1, Tbx6, Pax3, Vegfr2, Sox2* and *Sox10* in the subpopulations of Msgn1-repV cultures stained for PDGFRα. Cultures were differentiated in RDL medium for 3 (d3) and 4 (d4) days and sorted by FACS based on the expression of Msgn1-repV (M) and PDGFRα (Pα), as shown in (A), and compared to undifferentiated ES cells. Values are normalized to the M^-^Pα^-^ d3 condition. Mean+/- s.d. (D) qPCR analysis of the expression of anterior PSM/somitic markers *Pax3, Meox1/2, Tbx18, Uncx* and *Foxc2* in the subpopulations of Msgn1-repV cultures double stained for PDGFRα and VEGFR2. Cultures were differentiated in RDL medium for 4 (d4) days and sorted by FACS based on the expression of Msgn1-repV (M), PDGFRα (Pα) and VEGFR2 (V), as shown in (B), and compared to undifferentiated ES cells. Values are normalized to the triple negative population M^-^ Pα^-^V^-^ condition. Mean+/- s.d. (E) Representative FACS analysis of Pax3-GFP reporter cell cultures differentiated *in vitro* in (left) control FBS 15% (F15), in RDL medium without (center) or with pre-differentiation in NK1 medium (NK1- RDL, right) for 6 days and labeled with anti-PDGFRα or anti-VEGFR2 antibodies.

To further characterize these cell populations, we analyzed by qPCR the expression profile of various lineage markers in different fractions of differentiated Msgn1-repV^+^ cells sorted based on Venus and PDGFRα expression (as indicated in Figure 6A). We found that the Msgn1-repV^+^ PDGFRα^+^ (M+Pα+) fraction was strongly enriched for paraxial mesoderm markers such as *Msgn1, Tbx6, and Pax3* (Figure 6C). The Msgn1-repV^-^ PDGFRα+ (M-Pα+) fraction showed very low levels of *Tbx6* suggesting that it does not contain PSM cells (Figure 6C). These M-Pα+ cells expressed *Sox10* together with *Pax3* but lack *Sox2* expression suggesting that this fraction contains neural crest cells (Figure 6C). The Msgn1-repV subpopulations were further analyzed by qPCR following combined PDGFRα and VEGFR2 staining. While the Msgn1-repV^+^ PDGFRα+ VEGFR2^-^ (M+Pα+V-) population expressed the highest levels of anterior PSM/somitic markers, the triple positive M+Pα+V+ population also expressed a significant level of anterior PSM/somitic markers (Figure 6D). Interestingly, among the Msgn1-repV^-^ subpopulations, the PDGFRα^+^ VEGFR2^-^ (M-Pα+V-) also expressed comparable level of anterior PSM/somitic markers, suggesting that by day 4 a fraction of cells transited to anterior PSM/ somitic fate and downregulated the Msgn1-repV reporter. Together, our results indicate that the posterior PSM-like cells differentiated *in vitro* express PDGFRα, but they also argue for the lack of specificity toward the paraxial mesoderm lineage of the anti-PDGFRα antibody, even when combined with the VEGFR2 antibody.

To further establish the specificity or lack thereof of the PDGFRα and VEGFR2 surface markers in identifying somitic mesoderm, the Pax3-GFP mES line was differentiated for 6 days in RDL medium and analyzed for PDGFRα and VEGFR2 surface expression. Pax3-GFP^+^ cells differentiated with or without pre-differentiation in NK1 medium (Chal et al, 2015) were essentially negative for VEGFR2 while the majority were PDGFRα-positive (Figure 6E). Nevertheless, Pax3-GFP^+^ PDGFRα^-^ accounted for about 15 to 35% of the total Pax3-GFP^+^ population. Moreover, staining for CXCR4 showed that Pax3-GFP^+^ are negative for CXCR4 (Figure S2). Altogether, this suggest that PDGFRα, VEGFR2 and CXCR4 surface expression does not fully capture Paraxial mesoderm identity.

### Transcriptomic analysis of the differentiated Msgn1-repV and Pax3-GFP-positive cells

We next compared the identity of the transcriptome of the Msgn1-repV and Pax3-GFP-positive cells differentiated *in vitro* to their *in vivo* counterpart. Microarrays were generated for Msgn1-repV^+^ and Pax3-GFP^+^ cells sorted by FACS after 3, 4 or 5 days of differentiation in serum-containing RDL and CDL media. Their gene signatures were compared to that of the anterior and posterior PSM transcriptional domains *in vivo* (Chal et al., 2015) and with the neural tube array series to examine tissue specificity (Figure 7, Table S2, S5). As the differentiating ES cells transited from a Msgn1^+^ to a Pax3^+^ stage, they down-regulated a large number of posterior-specific PSM genes including *Dusp4, Rspo3, Evx1* and *Fgf8* as observed *in vivo* (Figure 7A, C). Interestingly, while many posterior PSM signature genes where also shared with the posterior-most neural tube, such was not the case for the anterior PSM signature genes. In parallel, progressive activation of a large fraction of the anterior PSM specific signature genes including *Ripply2, Mesp2, Nkx3-1, Tbx18*, was observed during *in vitro* differentiation (Figure 7B, D). Thus, *in vitro* differentiation of the Msgn1-repV and the Pax3-GFP reporter ES cells in the presence of Rspo3 and BMP inhibitors appears to recapitulate the early differentiation stages of the paraxial mesoderm *in vivo*.

**Figure 7:**
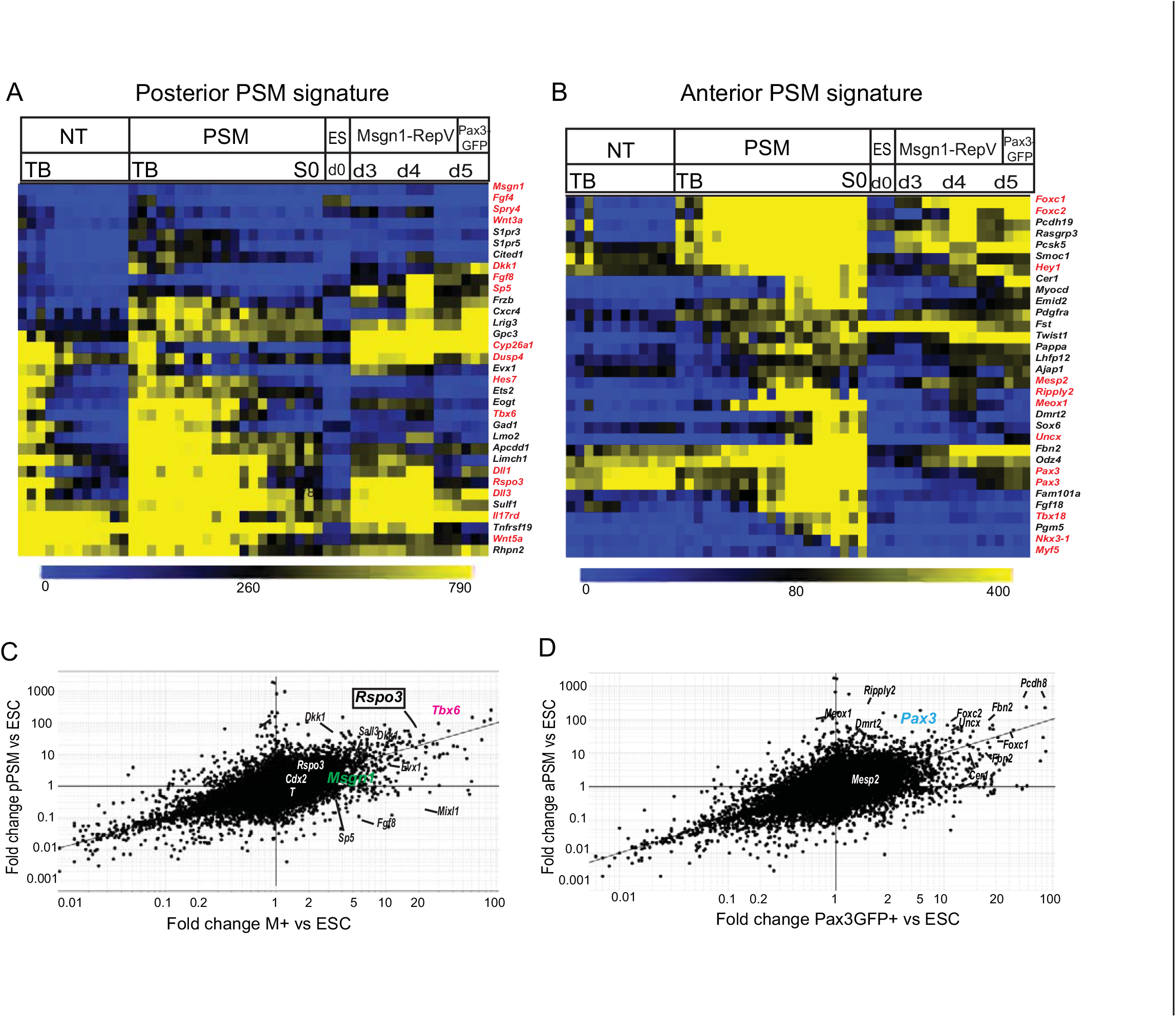
Gene signature comparison of mouse ES-derived PSM-like cells to PSM and NT tissues. (A, B) Heatmap of the expression levels of signature genes enriched in the posterior PSM (A) or anterior PSM (B) comparing the neural tube (NT) tissue series, the PSM tissue series, undifferentiated mES cells (ES), FACS-sorted Msgn1-repV^+^ and Pax3-GFP^+^ cells differentiated for 3 (d3), 4 (d4) and 5 (d5) days in RDL medium. Each row corresponds to a gene/probeset indicated on the right. Each column corresponds to an individual microarray. For each condition, biological triplicates are shown, except for NT (duplicates). Tissue series are oriented from the most posterior level (left, Tail bud (TB)) to more anterior levels (right). S0: forming somite. (C, D) Comparison of the transcriptional profiles of *in vitro* differentiated mouse ES-derived PSM-like cells to microdissected PSM tissue. Fold change versus fold change (FcFc) scatter plot comparing (C) FACS-sorted Msgn1-RepV^+^ (M^+^) cells differentiated for 4 days in RDL medium and posterior PSM (pPSM) or (D) FACS-sorted Pax3-GFP^+^ cells differentiated for 5 days in RDL medium and anterior PSM (aPSM) microarrays. Expression values are normalized to mouse ES cell microarray expression data sets. The expression ratio for each probe set and for each comparison is plotted on *x* axis and *y* axis, respectively (black dots, fold change, logarithmic scale). Known representative signature genes specific for the posterior PSM (C) or the anterior PSM (D) tissues are indicated.

### *In vivo* analysis of the myogenic potential of the ES-derived PSM-like cells

Trunk skeletal muscle tissue is generated exclusively from somitic paraxial mesoderm (Sambasivan et al., 2011). Properly specified PSM-like cells should therefore have the unique potential to generate skeletal muscle. We next analyzed the myogenic potential of the mouse PSM-like cells differentiated *in vitro* by transplanting them directly into injured adult muscles *in vivo*. Msgn1-repV^+^ and Pax3-GFP^+^ cells were differentiated for 3-4 days and 5-6 days in RDL medium respectively, and subsequently purified by FACS. Sorted cells were permanently labeled with a lentivirus driving ubiquitous expression of GFP in order to follow their fate after they are grafted *in vivo*. 50,000-100,000 labeled Msgn1-repV^+^ or Pax3-GFP^+^ cells were transplanted into cardiotoxin-injured *tibialis anterior* muscle of adult Rag2^-/-^ γc^-/-^ mice (Figure 8A). Freshly isolated, GFP-labelled adult satellite cells were also used as a positive control. One month posttransplantation, Msgn1-repV^+^ and Pax3-GFP^+^ donor cells reconstituted large areas filled with small, poorly organized, striated dystrophin-positive muscle fibers as well as occasionally other derivatives such as fibroblasts, chondrogenic nodules or epithelial cells forming large cysts (Figure 8B, C and data not shown). GFP-labeled muscle fibers expressed embryonic, slow or perinatal/fast isoforms of myosin heavy chain (MyHC) indicating that they span a large array of myogenic differentiation stages (Figure 8D). Thus, when transplanted *in vivo* into adult injured muscles, PSM-like cells derived *in vitro* from ES cells are able to continue their differentiation toward the myogenic lineage.

**Figure 8:**
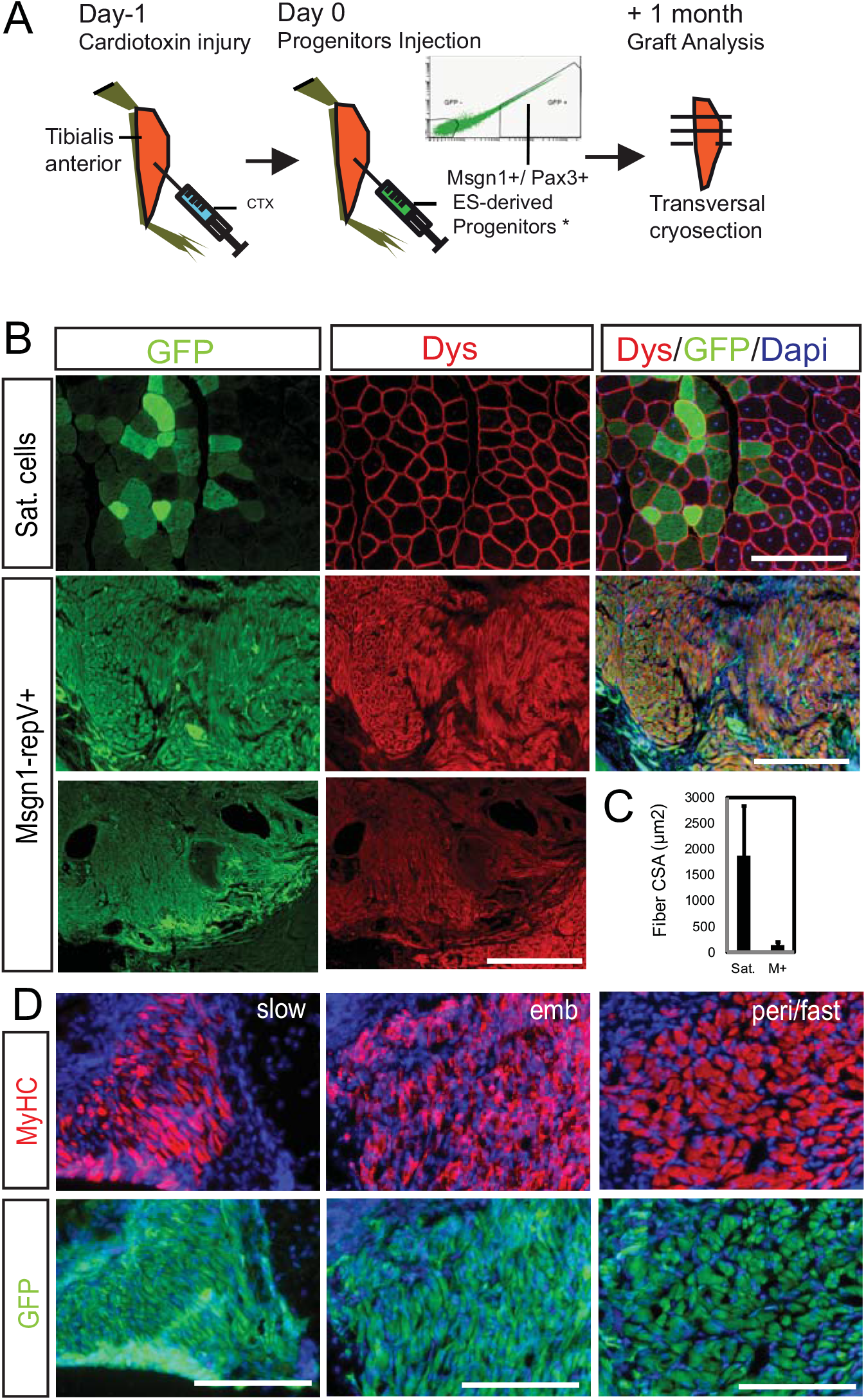
Transplantation of mouse ES-derived PSM-like cells leads to muscle formation *in vivo*. (A) Experimental scheme of the transplantation protocol of mouse ES-derived PSM-like cells into in the cardiotoxin-injured *tibialis anterior* of an adult Rag2^-/-^ γc^-/-^ mouse. (B) Representative transverse sections of mouse *tibialis anterior* muscles transplanted according to (A) with 30,000 satellite cells or with 50,000-100,000 FACS-sorted Msgn1-repV^+^ cells. Cells were permanently labeled with a GFP-expressing vector prior to injection. Transplanted cells’ contribution to muscle is visualized with an anti-GFP antibody (green) and individual fibers with an anti-dystrophin (Dys, red) antibody. Scale bar, 100μm, (bottom row) 400 μm. (C) Histogram showing the quantification of GFP^+^ fibers cross sectional area (CSA) generated from transplanted satellite cells or Msgn1-repV^+^ (M^+^) cells, respectively. Mean+/- s.d. (D) Representative transverse sections of mouse *tibialis anterior* muscles transplanted according to (A) with FACS-sorted Msgn1-repV^+^ cells. One month post-transplantation, fibers derived from the PSM-like cells (green) were positive for slow MyHC, embryonic MyHC and perinatal/fast MyHC (red) as detected by immunohistochemistry. Nuclei are counterstained (blue). Scale bar, 100μm.

### Differentiated PSM-like cells can recapitulate skeletal myogenesis *in vitro*

We next sought to define conditions in which the anterior PSM-like cells differentiated from ES cells could be reproducibly induced to generate skeletal muscle *in vitro*. In the embryo, FGF and Wnt signaling are downregulated in the anterior PSM prior to activation of the myogenic program (Aulehla and Pourquie, 2009). We therefore first differentiated ES cells in RDL medium for 4-6 days and then cultured cells for 2 days in a medium (thereafter referred as PDL) lacking the Wnt activator and containing the FGF inhibitor PD173074, while maintaining the BMP inhibition with LDN193189. Subsequently, cells were transferred to a myogenic differentiation medium containing 2% horse serum, in which they were maintained for up to 2 months. To monitor the generation of skeletal muscle cells, we used the Myog-repV ES reporter line described in (Chal et al., 2015). Myog-repV^+^ myocytes harboring a single centrally located nucleus, strongly resembling early myotome cells produced during primary myogenesis, were visible after 1 week in culture in these conditions (Figure 9A). We found that predifferentiating the mouse ES cell for 2 days in NK1 medium prior to exposure to RDL medium led to a more robust and homogeneous activation of the Myog-repV reporter in culture. Futhermore, substituting R-spondin3 by the GSK3β inhibitor and Wnt activator CHIR 99021 also led to efficient generation of Myog-repV^+^ myocytes. These myocytes were not observed when cells were maintained in RDL (or CDL) after day 6 or differentiated in base FBS 15% media (data not shown). The number of myocytes steadily increased over time progressively covering the entire surface of the wells (Figure 9D). Large numbers of elongated slow and perinatal/fast Myosin Heavy chain (MyHC) positive fibers progressively appeared from the Myog-repV^+^ myocytes cells (Figure 9B,C). While immature myogenic cells expressed MyoD and a limited subset also Myf5, myonuclei found in myotubes were positive for MyoD and Myogenin (Figure 9E and data not shown). By 2 weeks of differentiation, myotubes formed by fusion of myocytes which aligned spontaneously in culture, leading by three weeks *in vitro* to a large number of multinucleated Fast-MyHC^+^ fibers (Figure 9F-H) containing up to 50-100 myonuclei each, as seen in perinatal fibers *in vivo* (White et al., 2010) (Figure 9 G; data not shown). Moreover, larger fibers expressed Dystrophin in their sub-sarcolemmal domain (Figure 9I), and in some instance exhibited a single cluster of Acetylcholine receptors in equatorial position, as detected by fluorescent bungarotoxin (Figure 9J). Strikingly, the muscle fibers differentiated in culture for 3 to 5 weeks extended over several millimeters (Figure 9F-H) and reached a density of up to 80 fibers per square millimeter (data not shown). Around 10 to 15,000 such multinucleated muscle fibers could be obtained in wells initially seeded with 20 to 30,000 ES cells. Immunolabeling with anti-perinatal MyHC antibody showed that the generated fibers exhibit highly organized striation as expected from mature muscle fibers (Figure 9H). Many of the differentiated fibers contracted spontaneously indicating that the sarcomeric organization of the fibers is functional (Movie S1, S2). The dimensions of these fibers were between 1-3 millimeters in length and 10-20 micrometers in width, with a sarcomeric length around 2.5 micrometers, which are values similar to those of mouse post-natal fibers (White et al., 2010) (Figure 9L,M). Fibers were individually surrounded by a continuous basal lamina visualized by laminin deposits (Figure 9K). In long term cultures, muscle fibers were frequently found to develop over an epithelial-like sheet of cells (data not shown). Together, these data suggest that our *in vitro* culture conditions are able to recapitulate a myogenic differentiation sequence resembling that observed during normal embryogenesis in the mouse.

**Figure 9:**
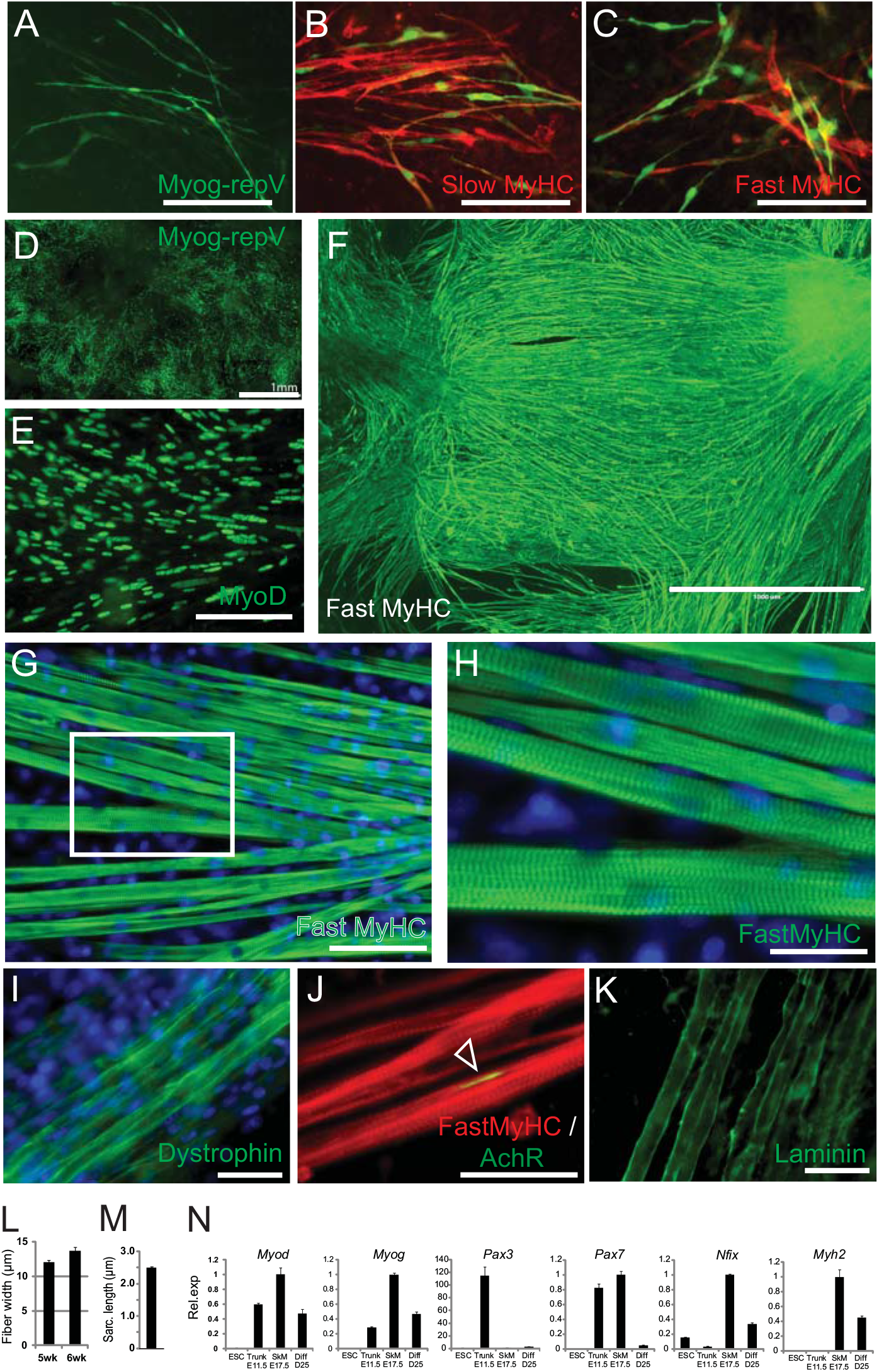
Differentiation of mouse ES-derived PSM-like cells *in vitro* recapitulates skeletal myogenesis. (A) Myog-repV cells differentiated for 8 days *in vitro* labeled with an anti-GFP antibody. Note the centrally located nuclei in elongated cells. Scale bar, 100μm (B, C) Immunostaining for slow MyHC (B, red) and peri/fast MyHC (C, red) on Myog-repV^+^ cells (green) differentiated for 10 days in vitro. Scale bar is 100μm. (D) Composite image of a large field of MyogRepV^+^ after 8 days in culture stained with an anti-GFP antibody. Scale bar, 1mm. (E) Representative mouse ES-derived myogenic culture obtained after 2 weeks according to the RDL protocol and stained for MyoD. Note the aligned MyoD^+^ myonuclei in nascent myotubes. Scale bar, 100μm (F) Large field of mature, millimeter-long myofibers stained for Fast MyHC. Mouse Pax7-GFP cells were differentiated in vitro for 3 weeks. Scale bar, 1mm. (G-H) Representative multinucleated myofibers differentiating from ES cells after 4 weeks in vitro. The culture was stained for Fast MyHC (green), nuclei are counterstained in blue. (G) Low magnification. Scale bar, 100 μm. (H) Higher magnification of the area shown in G (white boxe). Scale bar, 25 μm. Note the highly organized myofibrils and sarcomeric repeats. (I-K) Characterization of the skeletal muscle fibers generated *in vitro* from mouse ES cells. Fibers were stained for Dystrophin (I, nuclei counterstained in blue), for Acetylcholine Receptor (with Bungarotoxin) and Fast MyHC (J) and for Laminin (K), respectively. Scale bar, 50 μm. (L) Quantification of the width of mouse ES-derived myofibers at 5 or 6 weeks in culture. Mean+/- s.d. (M) Quantification of sarcomeric repeat length along myofibrils of mature muscle fibers differentiated from mouse ES-derived PSM-like cells *in vitro*. Mean+/- s.d. (N) qPCR analysis of the expression level of *MyoD, Myog, Pax3, Pax7, Nfix* and *Myh2* between ES cells, E11.5 embryonic trunk muscles, E17.5 fetal back muscles and mouse ES-derived PSM-like culture differentiated for 25 days. Values are normalized to E17.5 muscle sample. Mean+/- s.d.

We next used qPCR to analyze the expression level of a set of myogenic markers in 3-week old cultures differentiated in the conditions described above. These cultures were compared with undifferentiated ES cells, with E11.5 trunk muscles (containing primary myofibers) and E17.5 back muscles (in which foetal myogenesis is ongoing) (Figure 9N). While expression of the myogenic factors *MyoD* and *Myogenin* was clearly observed in the three muscle-containing samples, *Pax3* was only detected in the E11.5 sample. In contrast, *Pax7* was robustly expressed in both muscle samples and at a lower level in the differentiated ES myogenic cultures. The marker of fetal muscle fibers *Nfix* (Messina et al., 2010) and the *Myh2* (Fast2A MyHC) were not observed in E11.5 trunk muscles but were significantly enriched both in the E17.5 muscle sample and in the 25-day old cultures (Figure 9N). Thus, both primary and secondary (fetal) skeletal myogenesis can be recapitulated *in vitro* ultimately resulting in the production of striated contractile muscle fibers exhibiting a phenotype similar to early post-natal fibers.

### Production of ES-derived Pax7-positive myogenic precursor cells *in vitro*

During embryogenesis, muscle fibers are generated by proliferating progenitors expressing Pax7 and/or Pax3 that are located in between the growing fibers and that ultimately become located under the fiber basal lamina during late fetal stage to become satellite cells. We examined the presence of such Pax7^+^ progenitors in our long term cultures of differentiated ES cells described above. By two to three weeks of differentiation, large streams of Pax7^+^ cells were observed in the cultures and interspersed with newly formed myocytes suggesting that these correspond to myogenic progenitors (Figure 10A). Between three and four weeks, these populations quickly resolved to generate skeletal myocytes and aligned muscle fibers. By 4 weeks of differentiation and onward, the number of Pax7^+^ cells in culture decreased drastically. Nevertheless, a small population of Pax7^+^ was found in close association to large myofibers, a topography strikingly reminiscent of the *in vivo* situation where adult Pax7^+^ satellite cells are found in contact with fully differentiated muscle fibers (Yin et al., 2013) (Figure 10B, C). A fraction of Pax7^+^ progenitors become quiescent as suggested by the loss of Ki67 expression (Figure 10D-F, H). Thus our experiments show that differentiation of Satellite-like Pax7-positive myogenic progenitor cells can be obtained *in vitro* using our differentiation strategy.

**Figure 10:**
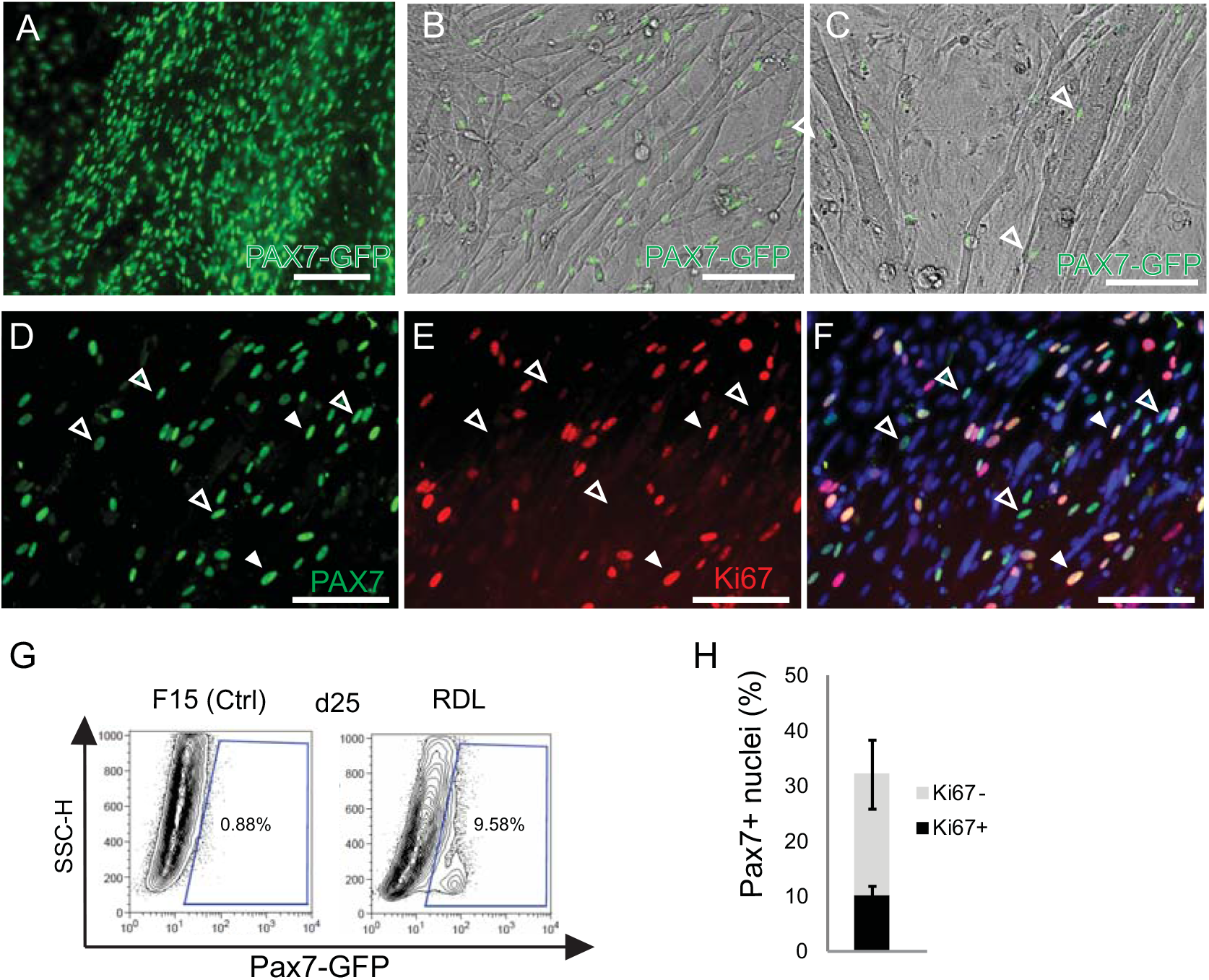
Differentiation of Pax7^+^ myogenic precursors from ES-derived PSM-like cells *in vitro*. (A-C) Live Pax7-GFP (green) expression in mouse ES cultures differentiated for 2 weeks in vitro (A) and at 4 weeks of differentiation (B, C). B and C are composite images merging the Pax7-GFP (green) signal and the corresponding transmitted light image (grey) highlighting myofibers edges. Note the juxtaposition of Pax7^+^ cells to the mature myofibers in 4 week-old cultures. Scale bar, 100μm. (D-F) Comparison of Pax7 (D, green) and Ki67 (E, red) immunolocalization in 2.5 week-old cultures. F corresponds to a merge of D and E with a nucleus counterstaining (blue). Double Pax7^+^/ Ki67^+^ nuclei (Full arrowheads) and single Pax7^+^/ Ki67^-^ nuclei (hollow arrowheads) are shown. Scale bar, 50 μm. (G) Comparison of the proportion of Pax7-GFP^+^ cells (blue gate) in 3.5 week -old mouse ES cultures differentiated according to myogenic induction protocol described or in FBS 15% control media (F15) and analyzed by flow cytometry. SSC-H: side scatter height. (H) Histogram showing the relative proportion of double Pax7^+^/ Ki67^+^ and single Pax7^+^ /Ki67^-^ nuclei in 2.5 week-old cultures as shown in (D-F). Mean +/- s.d.

## Discussion

Here, we describe a novel protocol recapitulating developmental cues *in vitro* to promote highly efficient myogenic differentiation of mouse ES cells. This protocol, which contains serum, allows for more effective differentiation and maturation of striated myofibers than the chemically defined protocols recently published (Chal et al., 2015; Shelton et al., 2014). Thus it might provide a useful tool to study aspects of myogenesis otherwise difficult to study *in vivo* such as myoblast fusion, myofibrillogenesis or satellite cell maturation. Activation of Wnt/β-catenin signaling combined to BMP inhibition leads to efficient differentiation of ES cells toward a posterior PSM identity. Further differentiation of these cells leads to the differentiation of Pax3-positive cells exhibiting characteristics of the anterior PSM fate. We also identified culture conditions allowing these PSM precursors to differentiate into Myogenin-positive mononucleated myotubes after 7 days. In long term culture, these cells produce large numbers of muscle fibers and their associated proliferating Pax7-positive precursors.

When allowed to differentiate in various conditions, embryoid bodies derived from mouse or human embryonic stem cells can yield a wide variety of derivatives from the three germ layers including paraxial mesoderm specific derivatives (Awaya et al., 2012; Braun and Arnold, 1994; Chang et al., 2009; Mizuno et al., 2010; Rohwedel et al., 1994). However, these approaches yield highly mixed populations and differentiation of mature derivatives such as skeletal muscle is poorly efficient and reproducible. Alternatively, differentiation of mouse and human pluripotent cells in monolayers combined with various treatments to modulate the Activin, BMP, and Wnt pathways has been reported to be able to induce a paraxial mesoderm fate (Sakurai et al., 2009; Sakurai et al., 2012; Tanaka et al., 2009). However, the methods reported in these studies lead to limited production of paraxial mesoderm precursors which need to be enriched by FACS-sorting to generate limited amount of paraxial mesoderm derivatives such as muscle or cartilage. Furthermore, these studies relied on the expression of PDGFRα to identify paraxial mesoderm cells. Our data show that whereas the Msgn1^+^ population of differentiated ES cells is recognized by the antibody against PDGFRα, this antibody also recognizes many other non-paraxial mesoderm cell populations. Hence, even in combination with the anti-VEGFR2 antibody, the anti-PDGFRα antibody is not specific for the paraxial mesoderm lineage and thus cannot be used as the identification criterion for this lineage. This is also supported by lineage tracing experiments in the mouse embryo (Ding et al., 2013; Ema et al., 2006; Motoike et al., 2003). Most studies published so far on the differentiation of paraxial mesoderm cells and their derivatives from ES cells, have relied on this identification method (Chan et al., 2016; Darabi et al., 2008; Filareto et al., 2012; Hwang et al., 2014; Magli et al., 2013; Sakurai et al., 2006; Sakurai et al., 2008; Sakurai et al., 2012; Tanaka et al., 2009) and thus the populations described as paraxial mesoderm likely correspond to a mixture of fates.

Using the Msgn1-repV and Pax3-GFP lines, we demonstrate that activation of the Wnt/β-catenin pathway with Rspo3 or CHIR-99021, can induce up to 70-90% of cells to activate the Msgn1-RepV reporter, which identifies a posterior PSM fate. Such a requirement of Wnt/ β-catenin signaling for paraxial mesoderm differentiation has been well established both *in vivo* in mouse mutants for the Wnt pathway (Dunty et al., 2008; Galceran et al., 2004; Takada et al., 1994; Yamaguchi et al., 1999), and *in vitro* in differentiating ES cell cultures (Borchin et al., 2013; Mendjan et al., 2014; Shelton et al., 2014). We further demonstrate that while Wnt activation alone is sufficient to activate expression of Msgn1 in mouse and human pluripotent cells, these cells start to express Bmp4 and progressively drift to a lateral plate fate, expressing markers such as *Foxf1* or *Hand1* and 2. *In vivo*, Bmp4 has been shown to be able to divert early paraxial mesoderm progenitors toward a lateral plate fate, consistent with its well described ventralizing activity on the developing mesoderm (Tonegawa et al., 1997). When mouse ES or human iPS cells were treated with Rspo3 or CHIR-99021 alone, we noticed an activation of *Bmp4* in the Venus -positive cells together with expression of lateral plate markers such as *Foxf1*. This raised the possibility that early Msgn1-repV cells exhibit a mixed unresolved paraxial mesoderm/lateral plate identity. Alternatively, some cells might have transiently expressed *Msgn1*, and retained the stable fluorescent Venus, while subsequently differentiating into lateral plate under the influence of Bmp4, whereas other Msgn1^+^ cells retained a paraxial identity thus generating an heterogenous population of Msgn1-repV^+^ cells composed of a mixture of lateral plate and paraxial mesoderm cells. Since Bmp4 has been shown to activate its own expression (Adelman et al., 2002; Blitz et al., 2000; Rojas et al., 2005; Schuler-Metz et al., 2000), we treated the cells with BMP inhibitors such as Noggin to block this endogenous BMP production. This led to an active downregulation of *Bmp4* and to the dose-dependent induction of a paraxial mesoderm fate. Interestingly, we observed significant differences towards induction of Pax3-positive cells *in vitro* between different BMP inhibitors. Whereas Noggin was able to efficiently contribute to induction of the posterior Msgn1-positive fate, only LDN-193189 (which inhibits Bmp signaling more strongly than Noggin) was found to be efficient for generating Pax3-positive cells *in vitro*. Alternatively, Ldn has been shown to also inhibit other kinases than the BMP receptors, such as FGFR1 (Vogt, 2011). These off-target effects could also account for the improved efficiency of Ldn on Pax3 induction. Together, our data demonstrate that active Wnt signaling together with BMP inhibition is required to differentiate ES cells toward the paraxial mesoderm lineage. BMP modulation as also been shown to be important for intermediate mesoderm and chondrogenic mesoderm induction (Craft et al., 2013; Craft et al., 2015; Morizane et al., 2015; Tanaka et al., 2009; Umeda et al., 2012; Zhao et al., 2014). These requirements are clearly different from those necessary to produce other mesodermal cell types such as hematopoietic and cardiovascular lineages where the BMP and Wnt pathway need to be activated simultaneously, followed by a subsequent Wnt signaling inhibition step required to induce differentiation of the cardiomyocyte lineage (Laflamme et al., 2007; Lian et al., 2012; Naito et al., 2006; Nostro et al., 2008; Ueno et al., 2007; Yang et al., 2008; Zhang et al., 2008).

Finally, we generated a microarray time series from fragments of microdissected embryonic neural tube and compared it to the PSM series (Chal et al., 2015). Clustering analysis of these microarrays revealed that unlike the PSM, the early neural tube differentiation is a progressive process with no abrupt transcriptional fate specification, suggesting that PSM and neural tube differentiation are controlled by distinct mechanisms (Oginuma et al., 2017; Olivera-Martinez et al., 2014; Ozbudak et al., 2010). Interestingly, we found that the gene signature the posterior-most neural tube microdissected fragment was largely overlapping with the Tail bud fragment of the PSM series, supporting the idea that the progenitors of both lineages share a common program/origin (Henrique et al., 2015).

## Acknowledgments

We thank Christopher Henderson for critical reading of the manuscript. We are grateful to Jennifer Pace and Tania Knauer-Meyer for their help; Laurent Bianchetti for Bioinformatic support. We thank Claudine Ebel from the Cytometry Facility at IGBMC and the IGBMC Microarray Facility for assistance. We thank the Pasteur Institute Cytometry and Animal Facilities and the IGBMC Cell Culture Facility for assistance. This work was supported by an advanced grant from the European Research Council (ERC-2009-AdG 249931 to O.P.), by the FP7 EU grant Plurimes (agreement no. 602423) and by a strategic grant from the French Muscular Dystrophy Association (AFM-Téléthon) to O.P. Microarrays data reported in this paper have been deposited in GEO database under the accession number GSE39615 and pending.

## Contributions

J.C. designed and performed experiments, analyzed data and coordinated the project. Z.A.T. designed and performed experiments, generated and characterized the hMSGN1-Venus line with help from B.G. M.O. performed the neural tube microdissection series. B.G., A.M. and G.G. carried out most of the mouse ES cell differentiation and characterization experiments under J.C.′s supervision. B.G. performed the *in situ* hybridization screen under J.C.′s supervision. O.S. characterized the Myog-repV line. P.M. and O.T. helped with microarray data analysis. A.B. contributed experimentally to the early project. L.K. provided technical support. J.-M.G. generated reporter constructs. M.K. established the hPS culture, provided expertise and coordinated the project. B.G.-M. and S.T. provided expertise and performed transplantation. O.P. supervised the overall project. O.P. and J.C. performed the final data analysis and wrote the manuscript.

## Supplementary Materials

### Materials and Methods

#### Mouse ES cell culture and differentiation

##### Maintenance

Mouse ES lines Msgn1-repV, Pax3-GFP, Myog-repV and Pax7-GFP have been described previously (Chal et al., 2015). Undifferentiated mouse ES cells were cultured on feeders (mitomycin-C inactivated mouse embryonic fibroblasts) at 37 °C in 5% CO2, in a maintenance medium composed of DMEM supplemented with 15% fetal bovine serum (FBS, Millipore), penicillin, streptomycin, 2 mM L-glutamine, 0.1 mM nonessential amino acids, 0.1% β-mercaptoethanol and 1,500 U/mL LIF. Prior to the NK1 pre-differentiation step, cells were first passaged twice onto gelatin-coated, feeder-free culture plates in 2i+LIF maintenance medium (Ying et al., 2008).

##### Analysis of R-spondin3 and Noggin action on mouse ES cells differentiation

ES cells were trypsinized and plated at approximately 10-20,000 cells/ cm^2^ in gelatin-coated 24 well plate in a DMEM-based medium, with 15% FBS (DF15), supplemented or not with Rspo3 (Peprotech, R&D Biosystems), CHIRON99021 (Chir; Tocris, Stemgent), Dkk1 (R&D Biosystems), 0.5% DMSO (Sigma), Noggin (R&D Biosystems) and LDN-193189 (Ldn, Tocris, Stemgent) at various concentrations. Media was typically changed every second day. Experiments were done in biological triplicates.

##### Serum-based differentiation of mouse ES cells toward a PSM-like fate

ES cells cultured on Feeders (+LIF) were trypsinized and plated as single cells at at approximately 10-20,000 cells/ cm^2^ in gelatin-coated, feeder-free, 96, 24 or 6-well plates directly DMEM-based medium, with 15% FBS (DF15), supplemented with recombinant 10 ng/ml Rspo3 (Peprotech, R&D Biosystems), 0.5% DMSO (Sigma), and 0.1 μM LDN-193189 (Ldn, Tocris, Stemgent) for 2 d. Alternatively, Rspo3 was replaced with the GSK3-β inhibitor CHIRON99021 (Chir; Tocris, Stemgent) at 1-3 μM. At 2 days of differentiation, medium was changed to a DMEM-based medium, with reduced serum, typically 1% FBS, 14% KSR (DK14F1) supplemented with Rspo3 (or Chir), DMSO and Ldn (RDL or CDL media) as indicated above. Alternatively, ES cells cultured on gelatin (+2i + LIF) were trypsinized and plated at 10-20,000 cells /cm^2^ on gelatin-coated, feeder-free, 96, 24 and 6-well plates and pre-differentiated in serum-free N2B27 medium supplemented with 1% Knock-out Serum Replacement (KSR, Gibco), thereafter (NK1 medium) for 2 days. Cells were then changed to the RDL (or CDL) medium described above. Media were refreshed every 2 days until day 6. While both methods generated PSM-like cells, the NK1 predifferentiation protocol was however more robust in generating large amount of PSM-like progenitors and was used for most of the long term *in vitro* myogenic differentiation. PSM differentiation experiments were performed at least 30 times independently, on 4 different mouse ES cell lines.

##### Serum-based skeletal muscle differentiation of the mouse PSM-like cells

Following 6 days of differentiation into PSM-like cells, cultures were changed to a DMEM-based reduced serum medium (DK14F1) supplemented with 0.25 μM of the MEK inhibitor PD0325901 (Stemgent) and 0.1 μM Ldn (PdL medium) for 2 days. After day 8 of differentiation, cultures were changed to a DMEM-based medium with 2% Horse serum (HS2%). Media was changed every other day. Myogenic differentiation experiments were performed at least 20 times independently, on 4 different mouse ES cell lines.

##### Neural differentiation

Protocol to generate Pax3^+^ neural rosette was based on “dual-smad” inhibition method described previously (Chambers et al., 2009). Pax3-GFP mouse ES cells were seeded as single cells as for mesodermal differentiation but cultured instead in a DMEM-based mediu with 15% FBS and supplemented with 10μM SB431542 and 0.1 μM LDN-193189 (Tocris).

#### Human iPS cell culture and differentiation

##### Maintenance

Undifferentiated human PS cell lines hiPS11a and H9 hES cells were cultured on Matrigel (BD Biosciences)-coated dishes in mTeSR1 media (StemCell Technologies). Cells were passaged as aggregates. Lines were regularly confirmed to be mycoplasma-free using a VenorGEM detection kit (Sigma).

##### Serum-free PSM-like and skeletal muscle differentiation of the human iPS cells

Human iPS cells were differentiated essentially according to (Chal et al., 2016). Briefly, hPS colonies were dissociated with Accutase (StemCell Technologies) and plated as single cells on Matrigel -coated 24 and 12-well plates (approximately 15,000-18,000 cells/cm^2^) in mTeSR1 supplemented with ROCK inhibitor (Y-27632, Sigma) for one day. The medium was changed to a DMEM-based medium supplemented with Insulin-Transferrin-Selenium (ITS, Gibco), 3 μM CHIRON99021 (Axon MedChem, Tocris) and 0.5 μM LDN-193189 (Axon MedChem, Stemgent) (CL medium). At day 3, 20 ng/ml FGF-2 (R&D Systems) was added for additional 3 d. After 6 d of differentiation, cells were changed to a DMEM-based medium supplemented with 10 ng/ml HGF, 2 ng/ml IGF-1, 20 ng/ml FGF-2 (R&D Systems) and 0.5 μM LDN-193189. After day 8 of differentiation, cells were cultured in DMEM, 15% KSR, supplemented with 2 ng/ml IGF1 for 4 days, and then supplemented with both 10 ng/ml HGF and 2 ng/ml IGF-1 after day 12 of differentiation. Media was changed every day until day 12, and every second day thereafter. Differentiation experiments were performed at least 15 times independently, on 2 unrelated human iPS lines. All human iPS and ES cell experiments were done according to local regulations (IGBMC and Brigham and Women’s Hospital) and in agreement with national and international guidelines.

##### Human hMSGN1-Venus hiPS reporter line generation

The *MSGN1* locus of undifferentiated hiPS11a was targeted by the CRISPR-Cas9 method (Addgene, (Cong et al., 2013; Ran et al., 2013)). sgRNAs targeting the stop codon of *hMSGN1* exon1 and a corresponding targeting vector, *hMSGN1-2A-nls-Venus,* containing a ~1.5kb 5’-homology arm (HA) followed by a 2A-peptide sequence (for translational cleavage) and a nuclearly localized (nls) Venus (YFP), followed by a Neomycin selection cassette and a ~1kb 3’HA. Heterozygous integrations were selected and coding regions were sequenced to verify that they do not contain indels. Three clones were further validated for correct expression and differentiation.

##### Flow cytometry analysis

Cells were trypsinized and analyzed by flow cytometry on a FACS calibur (BD Biosciences) according to reporter expression. Gating was determined for each reporter line using corresponding undifferentiated culture as a baseline control. Data are represented as % of FP^+^ cells in the culture. **Sorting**. Msgn1-repV^+^ (M^+^) and Pax3-GFP^+^ (P^+^) cells were isolated by FACS from Msgn1-repV and Pax3-GFP cultures respectively differentiated for 4-6 days in RDL (or CDL) medium. Biological triplicates were generated. Gated fractions were sorted either on FACS Aria (BD Biosciences) or S3 cell sorter (Bio-Rad). hMSGN1-Venus^+^ were sorted at 4 days of differentiation. Data were further analyzed with FlowJo software. Sorted populations were either processed for microarray analysis, qRT-PCR or for transplantation experiments. **Immunophenotyping**. Cells were trypsinized, resuspended in blocking solution PBS- 5% FBS supplemented with 5μM Rock inhibitor (Y-27632, Sigma) and incubated for 10 min at 4C. Conjugated primary antibodies, mouse anti-PDGFRα (CD140a, clone APA5)-APC and mouse anti-VEGFR2 (Flk-1/KDR/CD309, clone Avas12a)-PE (eBioscience) were added at 1μg and 0.5μg per million of cells, respectively. Corresponding isotype staining controls were also ran in parallel. Cells were incubated for 20 mins at 4C. Next, cells were washed with PBS and resuspended in PBS-2% FBS and analyzed or sorted for qPCR analysis. Flow cytometry analyses were performed at least three times independently.

##### Quantitative RT-PCR

Total RNA was extracted from PSC cultures using Trizol (Invitrogen) or with the RNeasy microkit (Qiagen). RT-PCR was performed on 5 ng total RNA using QuantiFast SYBR Green RT-PCR Kit (Qiagen) and gene-specific primers (Primerbank) and run on a LightCycler 480II (Roche). β-actin was used as an internal control for mouse system, and GAPDH for the human system. For mouse tail/PSM reference, mouse E9.5 tails were microdissected posterior to the level of the forming somite (S0) and pooled for total RNA extraction. For embryonic and fetal muscle references, CD1 mouse E11.5 trunk muscles and mouse E17.5 back muscles were microdissected for total RNA extraction.

##### Microarrays generation

Generation of the mouse neural tube (NT) microarray series was done as described previously for the PSM microarray series (Chal et al., 2015). Briefly, CD1 mouse E9.5 embryos were microdissected into caudo-rostral serie of consecutive ~100 μm fragments. Two series of 6 fragments were generated, from the tail bud to the newly formed somite (S0) level (Figure. 1B). RNA from each fragment was extracted with Trizol (Invitrogen) and used to generate probes hybridized on GeneChip Mouse Genome 430 2.0 arrays, as described previously (Chal et al., 2015). Differentiatied mouse ES and human iPS cultures were dissociated, FACS-sorted for reporter expression and processed as described previously (Chal et al., 2015). Mouse ES cells samples were hybridized on GeneChip Mouse Genome 430 2.0 arrays, while hiPS samples were hybridized on GeneChip Human Genome U133 2.0 arrays (Affymetrix).

##### Microarray data analysis

Initial filtering and preprocessing, including background correction, quantile normalization and summarization, was performed using both RMA and MAS with the R Bioconductor package (R version 2.12.1, Bioconductor version 2.8). Expression sets were then filtered according to Calls information. Probe sets expression fold changes between conditions (biological duplicates or triplicates) were calculated using the “Comparative Marker Selection” module of GenePattern (Reich et al., 2006). Histogram expression profiles of gene probesets were generated from MAS values. Further analysis was performed using the Manteia database (Tassy and Pourquie, 2014). Hierarchical clustering was performed on Microarray RMA data. Clustering was computed with an Average linkage method and Euclidean distances. Both an Approximately unbiased (AU) and Bootstrap probability (BP) P values were calculated (pvclust, R package). Clusters with AU and BP P values > 0.95 are highlighted by color-coded boxes. Expression heatmaps were generated with TM4-MeV (Saeed et al., 2006).

##### Gene signature lists (GSL) method

A gene expression reference was created by using all microarrays of wild-type mouse tissues deposited in GEO corresponding to Affymetrix Mouse Genome 430 2.0 Arrays (GEO platform id GPL1261) as of 08-2011. 1320 microarrays data sets from 255 distinct experiments were downloaded using a Perl script (Bioperl). Normalization was done by calculating the mean values for each microarray. The median values for the distribution of those mean values across all microarrays were determined. This median was then used as a scaling factor for each value on each microarray. Once all microarrays had been normalized, the median expression value for each probeset was defined as the reference value for that particular probeset/gene. Gene signature list specific to one experimental condition was generated by normalizing the corresponding microarray data as done for the reference dataset. Signature gene for a given conditions were any probeset whose normalized expression value was 10 times higher than the corresponding reference value.

##### PSM-like cells preparation for transplantation into injured *tibialis anterior* (TA) muscle

Msgn1-repV or Pax3-GFP mouse ES cells were differentiated for 4-6 days and pretreated with ROCK inhibitor (Y-27632, Tocris) one day before being trypsinized. One day prior to injection, the *tibialis anterior* muscles of a cohort of Rag2^-/-^ γc^-/-^ mice (see below) were injected with 10μM Cardiotoxin (Sigma). Cell preparation was filtered through a 30μm mesh and reporter-positive cells were sorted using a FACS Aria II or a Moflow Astrios (BD Biosciences). Sorted cells were replated and permanently labeled by transduction overnight with a CAG-GFP lentivirus (MOI of 20-30). Cells were then washed several time and re-incubated for 1 hour in medium at 37C before preparation for transplantation (Gayraud-Morel et al., 2012). For control satellite cells transplantation, Pax7-GFP^+^ were freshly isolated from Pax7-GFP mouse and labeled with a CAG-GFP lentivirus as described (Gayraud-Morel et al., 2012). 50,000 to 100,000 cells were injected in the tibialis anterior (TA) muscles of 3-4 month-old Rag2^-/-^ γc^-/-^ male mice. Injections were done under general anesthesia. Grafted *tibialis anterior* muscles were collected 1-2 months post transplantation and processed for cryosection and immunohistochemistry (Gayraud-Morel et al., 2012). Transplantation experiments were done at least 3 times independently on animal cohorts of 3-5 animals. Experiments on mice were done according to local regulations (Institut Pasteur, IGBMC), in agreement with national and international guidelines.

##### *In situ* hybridization

Whole mount *in situ* hybridization was carried out as described (Henrique et al., 1995). Antisens mRNA probes for novel PSM markers identified by microarray and gene signature analysis were synthesized from mouse E.9.5 embryonic tail cDNA with sequence specific primers (Sigma, Table S8). Images were acquired on a Leica M125 stereomicroscope. For each probe, at least 3 embryos were hybridized.

##### Immunohistochemistry

For cell cultures, cultures plates were fixed in 4% formaldehyde for 20 mins at 25 °C or overnight at 4°C. Cultures were rinsed three times in PBS, followed by incubation in blocking buffer composed of Tris-buffered saline (TBS) supplemented with 1% FBS and 0.1% Triton X-100 for 30 mins. Primary antibodies were then diluted in blocking buffer and incubated overnight at 4 °C. The next day, cultures were washed three times with TBST (TBS supplemented with 0.5% Tween-20) and incubated with secondary antibodies (1:500) and counterstained with DAPI or Hoechst (1:1000) for at least 6 hours. Cultures were finally washed with TBST followed by PBS, before analysis.

For *tibialis anterior* (TA) muscle, dissected TA muscles were prepared for cryosection according to (Gayraud-Morel et al., 2012). 12 μm transversal serial sections were prepared and incubated overnight with primary antibodies in blocking buffer on a Sequenza rack system (Shandon). Following TBST washes, slides were incubated with secondary antibodies conjugated with AlexaFluor (Molecular probes) at 1:500 in blocking buffer, and DAPI or Hoechst (1:1000) were used as counterstain. Following TBST and PBS washes, slides were mounted with Prolong Gold Antifade (Molecular Probes). Primary antibodies used in this study were anti-GFP (Abcam; 1:300), anti-Tbx6 (Abcam; 1:400), anti-MyoD (5.8A, Santa Cruz; 1:200), anti-Myogenin (F5D, DSHB; 1:250), anti-Pax3 (DSHB; 1:250), anti-Pax7 (DSHB; 1:250), anti-Slow MyHC (NOQ7.5.4D, Sigma; 1:250), anti-Embryonic MyHC (F1.652, DSHB; 1:250), anti-Fast MyHC (MY-32, Sigma; 1:300), anti-dystrophin (Mandra1, Sigma; 1:400), anti-dystrophin (DY4/6D3, Leica; 1:200), anti-Laminin (Abcam; 1:800) and anti-Ki67 (Abcam; 1:300). AchR was detected with conjugated Bungarotoxin (Molecular Probes).

##### Quantifications

Myofiber cross sectional area (CSA) was measured from immunostained (Dystrophin) transverse section of engrafted TA using Fiji (Schindelin et al., 2012). Two independent engrafted muscles were analyzed for each conditions and the CSA of >150 individual fibers were measured per muscle. *In vitro* mouse ES-derived myofiber width was measured from immunostained (Fast MyHC) fibers in differentiated cultures using Fiji. For each condition, three fields of fibers were analyzed and the width of >70 individual fibers were measured per cultures. *In vitro* sarcomeric length was measured from immunostained (Fast MyHC) fibers in differentiated cultures using Fiji. Measurement were done along the main axis of the fiber, perpendicular to the striations. Fifteen individual mature fibers were identified and for each of them >20 consecutive sarcomeric units were measured. *In vitro* myogenic marker expression was quantified from immunostained cultures. For each condition, >5 myogenic fields per well were quantified. Results are expressed as a % of positive nuclei.

##### Image acquisition and processing

Live or fixed samples were acquired either on a Zeiss Axiovert or Evos FL for systems for transmitted light and/or fluorescent images. Images were processed with Adobe Photoshop and measurements were done with Fiji. For muscle contractions, movies were taken with a Leica DMRB microscope using a photometric FX camera and a 20X objective. Images were taken at 10Hz. Image sequences were processed in Fiji and saved in AVI format at real time speed.

##### Statistical analysis

For array data hierarchical clustering, Approximately unbiased (AU) and BP P values were both calculated (pvclust, R package). Clusters with AU and BP P > 0.95 were considered significant. For differential array expression data comparison, unpaired t-test P values were calculated using the Comparative Marker Selection module of GenePattern (Reich et al., 2006). Differences were considered significant for P values ≤ 0.05. For qRT-PCR data, unpaired Student t-test and twotailed P values were calculated. Differences were considered significant for P values ≤ 0.05.

## Supplementary Materials

**Figure S1:**
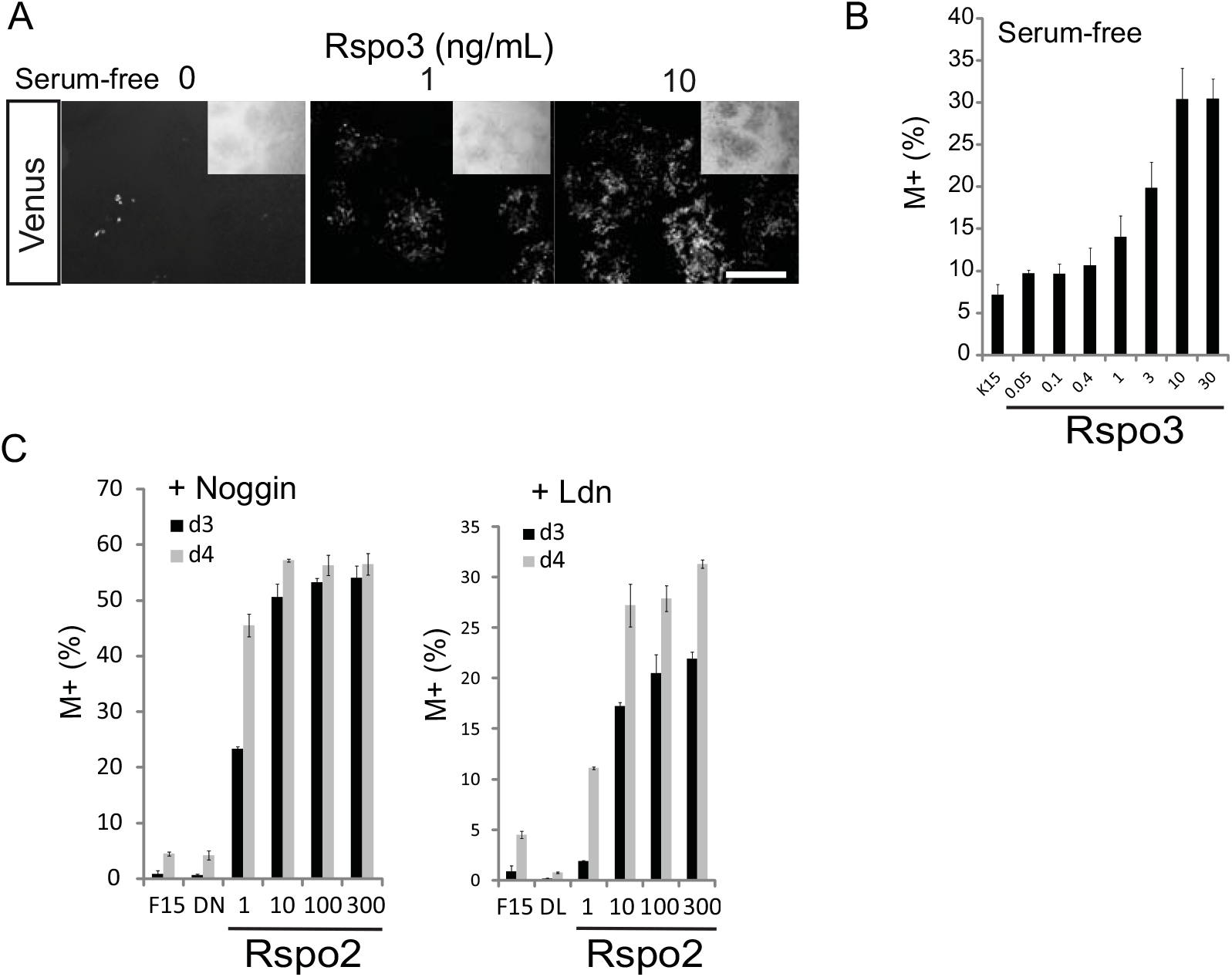
R-spondins induce the Msgn1-repV reporter in differentiating mouse ES cell cultures. (A) Activation of Msgn1-repV reporter with Rspo3 in serum-free conditions. Venus detection (white signal) of the reporter activation after 4 days of differentiation of Msgn1-repV ES cells, in a DMEM-based medium with 15% serum replacement (KSR) and supplemented or not with recombinant Rspo3. A transmitted light picture of each culture is shown in inset for each conditions. Scale bar, 200μm (B) Quantification of the proportion of Msgn1-repV -positive (M^+^) cells in cultures differentiated for 4 days in serum-free medium containing 15% KSR (K15) without or with various concentrations of Rspo3 (in ng/mL). Mean+/- s.d. (C) Induction kinetic of the Msgn1-repV reporter (M^+^) in mouse ES cell culture in presence of Rspo2. Cultures were differentiated for 3 (d3) or 4 (d4) days in either control DMEM-based media containing 15% FBS alone (F15), or supplementated with Dmso 0.5% and Noggin (DN, Left graph), or Ldn (DL, Right graph), and with various concentrations of Rspo2 (in ng/mL). Mean+/- s.d.

**Figure S2:**
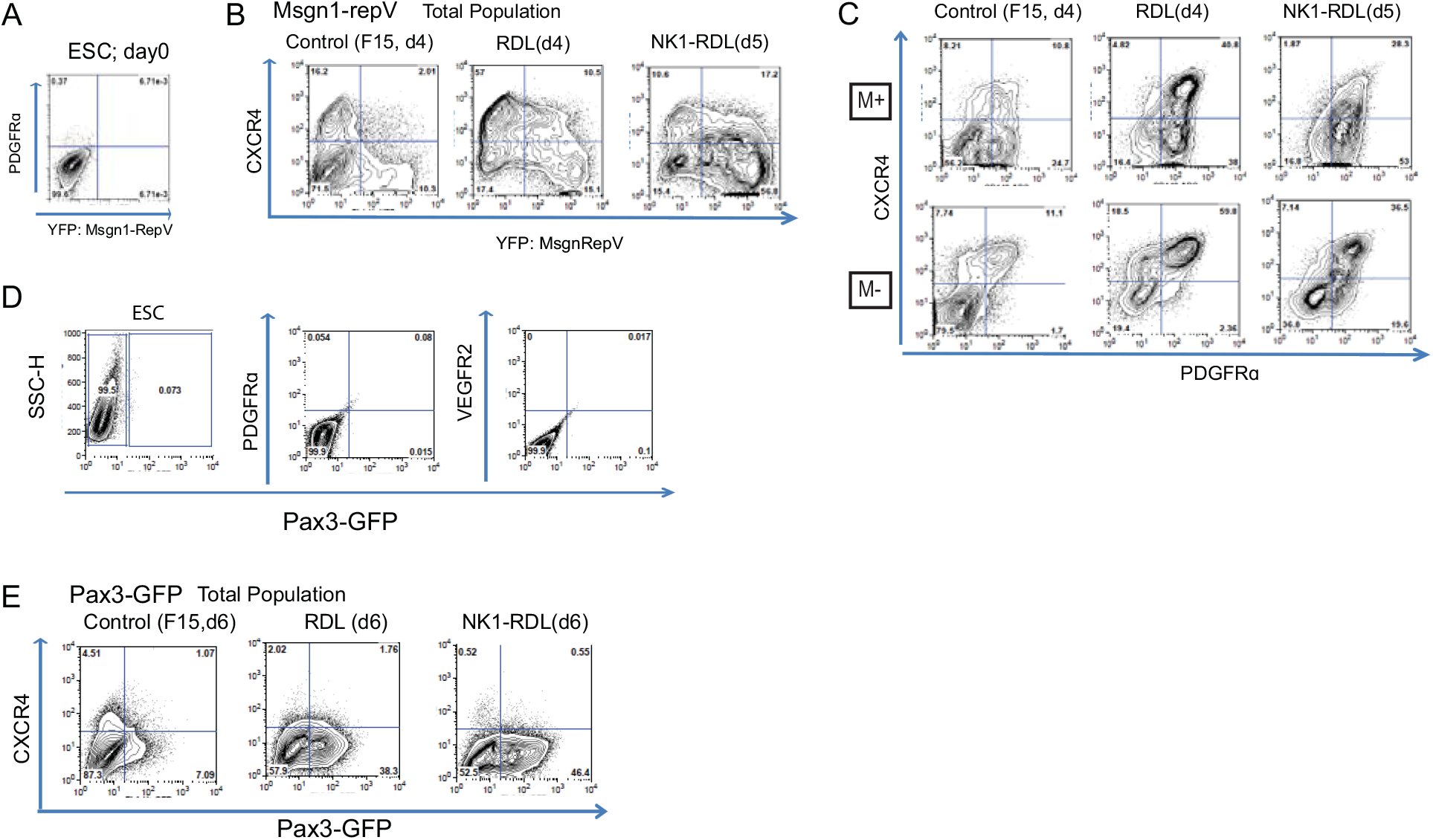
Immunophenotyping of mouse ES-derived PSM-like cells for PDGFRα, VEGFR2 and CXCR4. (A) Representative FACS analysis of undifferentiated (day0) mouse ES Msgn1-repV cells for PDGFRα expression. (B) Representative FACS analysis of Msgn1-repV reporter cell cultures differentiated *in vitro* in (left) control DMEM-based 15% FBS (F15), in RDL medium without (center) or with predifferentiation in NK1 medium (NK1- RDL, right) for 4-5 days and labeled with an anti-CXCR4 antibody. (C) Representative FACS analysis of PDGFRα and CXCR4 expression in the Msgn1-RepV-positive (M+) and -negative (M-) populations, differentiated in (left) control DMEM-based 15% FBS (F15), in RDL medium without (center) or with pre-differentiation in NK1 medium (NK1- RDL, right) for 4-5 days. (D) Representative FACS analysis of undifferentiated (day0) mouse ES Pax3-GFP reporter cell cultures for PDGFRα or forVEGFR2 expression. SSC-H: side scatter height. (E) Representative FACS analysis of mouse ES Pax3-GFP reporter cell cultures differentiated *in vitro* in (left) control DMEM-based 15% FBS (F15), in RDL medium without (center) or with predifferentiation in NK1 medium (NK1- RDL, right) for 6 days and labeled with an anti-CXCR4 antibody.

**Table S1: List of genes differentially expressed between posterior and anterior PSM transcriptional domains**. Expression intensity of each expressed probeset (RMA annotation) were compared between the posterior PSM domain (fragments 2 and 3 of PSM array series) and the anterior PSM domain (fragments 4, 5 and 6 of PSM array series). Comparison was done by Significance Analysis of Microarrays (SAM) analysis using MeV4.5 (TM4 Microarray Software suite). For each probeset, the corresponding gene and the relative expression fold change between the posterior and anterior PSM domains are shown. The False Discovery Rate was fixed at 0.525. Genes validated by *in situ* hybrization in this study are highlighted (yellow), or validated genes in the literature are also highlighted (orange).

**Table S2: List of signature genes for posterior and anterior PSM domains used for expression heatmaps**. For each marker gene, Affymetrix probeset identity is shown. Lists were assembled from this study and published literature.

**Table S3: Complete lists of signature genes for each Paraxial mesoderm and Neural tube domains (GSL method)**. Signature gene lists were generated by the GSL method (see Materials and Methods for details) for each paraxial mesoderm domains in the PSM array serie, namely Tail bud, Posterior PSM, Anterior PSM and Somite. The neural tube array series was arbitrarily subdivided into a posterior and anterior domains and signature gene lists were generated for both.

**Table S4: Venn diagram comparing the gene signatures list (GSL) of PSM versus Neural tube (NT) array series**. (left) Comparison between the core Neural tube GSL genes and the core PSM GSL genes. Core signature genes are genes found significantly upregulated in domain of a serie for a given tissue. (right) Comparison between all the signature Neural tube genes with all the signature PSM genes. For each subgroup, the corresponding list of signature genes is provided.

**Table S5: Lists of signature genes for mouse ES cells and differentiated FACS-sorted Msgn1-RepV-positive and Pax3-GFP-positive populations (GSL method)**. Signature gene lists were generated by the GSL method (see Materials and Methods for details) for undifferentiated mouse ES cells and differentiated populations (Msgn1-repV^+^, Pax3-GFP^+^) for 4 to 5 daysof differentiation in Rspo3/DMSO/Ldn (RDL) or Chir/DMSO/Ldn (RDL) medium. The top 400 probe sets are provided for each conditions. For each probe set, the corresponding gene and the relative expression fold change compared to the reference median value are shown.

**Table S6: List of genes differentially expressed in in 4 day-old hMSGN1-GFP-positive cells cultured in Chir/Ldn or Chir only media versus undifferentiated hiPSC**. (TabA) List of genes upregulated >2 folds in MSGN1-GFP+ cells in Chir/LdnL versus undifferentiated hiPSC. (TabB) List of genes upregulated >2 folds in MSGN1-GFP+ cells in Chir only versus undifferentiated hiPSC

**Table S7: List of genes differentially expressed in 4 day-old hMSGN1-GFP-positive cells cultured in Chir/Ldn versus Chir only media**. (TabA) List of genes upregulated >2 folds in hMSGN1-GFP+ cells in Chir/LdnL vs Chir only conditions. (TabB) List of genes upregulated >2 folds in MSGN1-GFP+ cells in Chir only vs Chir/Ldn conditions. For each probeset/gene Fold change (red) and False Discovery Rate (FDR(BH), yellow) are highlighted.

**Table S8: *In situ* probes information**

For each gene/probe, the ENSEMBL reference sequence number is provided. The primer couples sequences (F: forward, R: reverse) used to amplify the region of interest and the corresponding probe template size are provided.

**Movie S1: Spontaneous contractile activity of differentiated mouse ES-derived myofibers**. Mouse ES cells were differentiated in to PSM-like cells according to the RDL medium protocol and subsequently cultured in HS2% of additional weeks and imaged with transillumination. Real time speed.

**Movie S2: Spontaneous muscle bundle formation and contractile activity in adherent differentiated ES cultures**. Mouse ES cells were differentiated in to PSM-like cells according to the CDL medium protocol and subsequently cultured in HS2%. Large fiber bundle formed spontaneously *in vitro* anchoring only by extremities (not shown). Note synchroneous twitching of the bundle of aligned myofibers. Real time speed.

